# Spatiotemporal mapping of the contractile and adhesive forces sculpting early *C. elegans* embryos

**DOI:** 10.1101/2023.03.07.531437

**Authors:** Kazunori Yamamoto, Sacha Ichbiah, Matthieu Perez, Joana Borrego-Pinto, Fabrice Delbary, Nate Goehring, Hervé Turlier, Guillaume Charras

## Abstract

Embryo shape is determined by individual cell mechanics, intercellular interaction strength, and geometrical constraints. Models based on surface tensions at cell interfaces can predict 3D static cellular arrangements within aggregates. However, predicting the dynamics of such arrangements is challenging due to difficulties in measuring temporal changes in tensions. Here, we characterise the spatiotemporal changes in cellular tensions shaping the early nematode embryo using AFM, live microscopy, and tension inference. Using excoriated embryos, we validate a hybrid inference pipeline that calibrates relative inferred tensions temporally using cortical myosin enrichment and absolute tensions using AFM measurements. Applied to embryos within their native shell, we infer a spatiotemporal map of absolute tensions, revealing that ABa, ABp, and EMS compaction is driven by increased tension at free surfaces, while P_2_’s initial exclusion is due to high tension at intercellular contacts. We uncover a direct and non-affine contribution of cadherins to cell-cell contact tension, comparable to cadherins’ indirect contribution via actomyosin regulation.

**Highlights:**

- P lineage cells have lower cortical tensions than AB lineage cells
- Enrichment of Myosin-II at the cell cortex is a good predictor of cell-medium tension but is not sufficient to determine tension at cell-cell contacts.
- Myosin-informed tension inference allows determination of the spatiotemporal evolution of all surface tensions within the embryo.
- ABa, ABp, and EMS compact due to high tensions at their cell-medium interfaces compared to their cell-cell interfaces, while P_2_ is initially excluded due to high cell-cell contact tensions.
- Cadherins contribute directly in a non-linear way by reducing cell-cell contact tension by nearly 50%.

**Open Access:** For the purpose of Open Access, the author has applied a CC BY public copyright license to any Author Accepted Manuscript version arising from this submission.

## Introduction

Embryo shape is governed by cell mechanics, intercellular interaction strength, and geometric constraints. During development, cells continuously alter their mechanical properties, generate forces, and modulate interactions in response to developmental cues. These changes drive tissue morphogenesis. Our understanding of the forces shaping embryos has advanced significantly over the last decades, particularly in later developmental stages involving epithelial tissues, such as convergence and extension or gastrulation ^1–6^. Other studies have focused on early developmental stages where cell groups undergo slow, uniform mechanical changes ^7,8^. In all cases, the dynamics of mechanical changes were sufficiently slow to ensure a quasi-static evolution of cell mechanics. Early *C. elegans* morphogenesis, however, has been less studied, despite its widespread use as a model for genetics and development. Early cell cycles last only 10-20 minutes, with cell type specified from the 2-cell stage via asymmetric divisions ^9,10^, rotational movements ^11,12^, Wnt ^13^ and Notch signaling ^14,15^. Those processes are tightly controlled, ensuring an invariant development with minimal morphogenetic variation among embryos. Compared to other model systems, such as mouse ^8^ and human preimplantation embryos ^16^, *Drosophila* retina ^17,18^, ascidians ^18^ or frog gastrula ^19^, both morphogenesis and lineage specification in the *C. elegans* embryo are more rapid, which presents a true challenge for mechanical characterization and modeling ^20–24^.

Extensive knowledge has been gained on signaling and cell fate specification in early *C. elegans* embryos, but how molecular regulators control blastomere mechanics, shape and arrangement in the early embryo remains relatively underexplored. In many organisms, changes in cell shape and embryo morphology appear mostly regulated by differences in myosin enrichment at cell interfaces. For example, in the 8-cell stage mouse embryo, myosin increases at cell-medium surfaces and decreases at cell-cell interfaces, driving compaction ^8^. Similar myosin roles likely exist in *C. elegans*, where early divisions result in asymmetric myosin segregation across daughter cells ^9,10^. Cell adhesion proteins, such as cadherins, play an integral role in shaping cell aggregates through direct and indirect contributions to tension at intercellular contacts ^25–28^. In principle, cadherins might decrease interfacial tension directly through their contribution to adhesive energy, but in organisms like zebrafish ^29^ and mouse ^30^, this direct contribution is negligible compared to their indirect signalling effect. Indeed, cadherin binding decreases myosin at intercellular contacts, due to downregulation of RhoA after p120-catenin is recruited by the cadherin cytoplasmic domain ^31^. In *Drosophila*, cadherins also have a weak direct contribution coupled to a more significant indirect role ^32^. The relative importance of myosin activity and intercellular adhesion in surface tension control remains unclear in *C. elegans*, partly due to the challenge of characterizing the mechanics of the early embryo surrounded by an eggshell.

Recently, various computational models have been proposed to investigate the cell-specific mechanical changes driving *C. elegans* embryo morphogenesis. These include: (1) center-based models, which approximate cells as spheres interacting through central forces and enable the study of dynamic rearrangements across divisions but neglect surface forces that predominantly shape blastomeres^20,23^; (2) phase-field models, which provide a more precise representation of cell shape and mechanical interactions but require a tight balance of repulsive, tensile, and adhesive forces, making experimental calibration and validation challenging^21,22,24^; and (3) deformable cell models, which describe cells as interacting manifolds and necessitate numerous cell-specific parameters^33^ (such as repulsion, adhesion, viscosity, and contractility—many of which remain difficult to measure or calibrate directly from experiments).

Here, we adopt a foam-like model, an intermediate approach that has already demonstrated its relevance and efficiency to precisely describe cell shape and arrangement in early embryos ^8,16^. In this framework, cell-cell contacts are represented by a single interface with an effective surface tension that integrates contributions from both contractile and adhesive forces. This allows to characterise the spatiotemporal changes in cellular surface tensions that shape the early nematode embryo with a minimal number of relevant mechanical parameters. By combining direct AFM surface tension measurements of excoriated embryos with tension inference based on microscopy imaging, we show that the 4-cell stage arrangement in *C. elegans* arises from ABa, ABp, and EMS compaction due to an increase in tension at free surfaces, while P2 is initially excluded because of high intercellular tension at its contacts. Finally, we reveal that cadherins directly contribute to cell-cell interface tension, with a non-affine effect that is comparable in magnitude to their indirect contribution via actomyosin depletion. This direct contribution appears crucial in *C. elegans* for understanding how temporal changes in surface tension drive cell arrangement.

## Results

### Early *C. elegans* development involves dynamic, cell- and stage-specific changes in mechanics

Despite its small number of cells, the early *C. elegans* embryo acquires complex shapes that point to mechanical heterogeneity of cells and cell interfaces (**Fig 1A**). The one cell embryo undergoes an asymmetric division to give rise to AB and P_1_. AB then divides symmetrically into ABa and ABp, which are identified based on their position within the embryo; whereas P_1_ divides asymmetrically into EMS and P_2_. Together, ABa, ABp, and EMS form a compact aggregate, with P_2_ bulging out from this group (**Fig 1A, S1A**, 4-cell stage normalised time=0.1). Further divisions beyond the 4-cell stage give rise to complex non-planar 3D packing of the cells within the embryo but clear differences in contact angles between cells can nevertheless be observed. Overall, these observations imply dynamic and cell-specific regulation of mechanics within the embryo.

**Figure 1:**
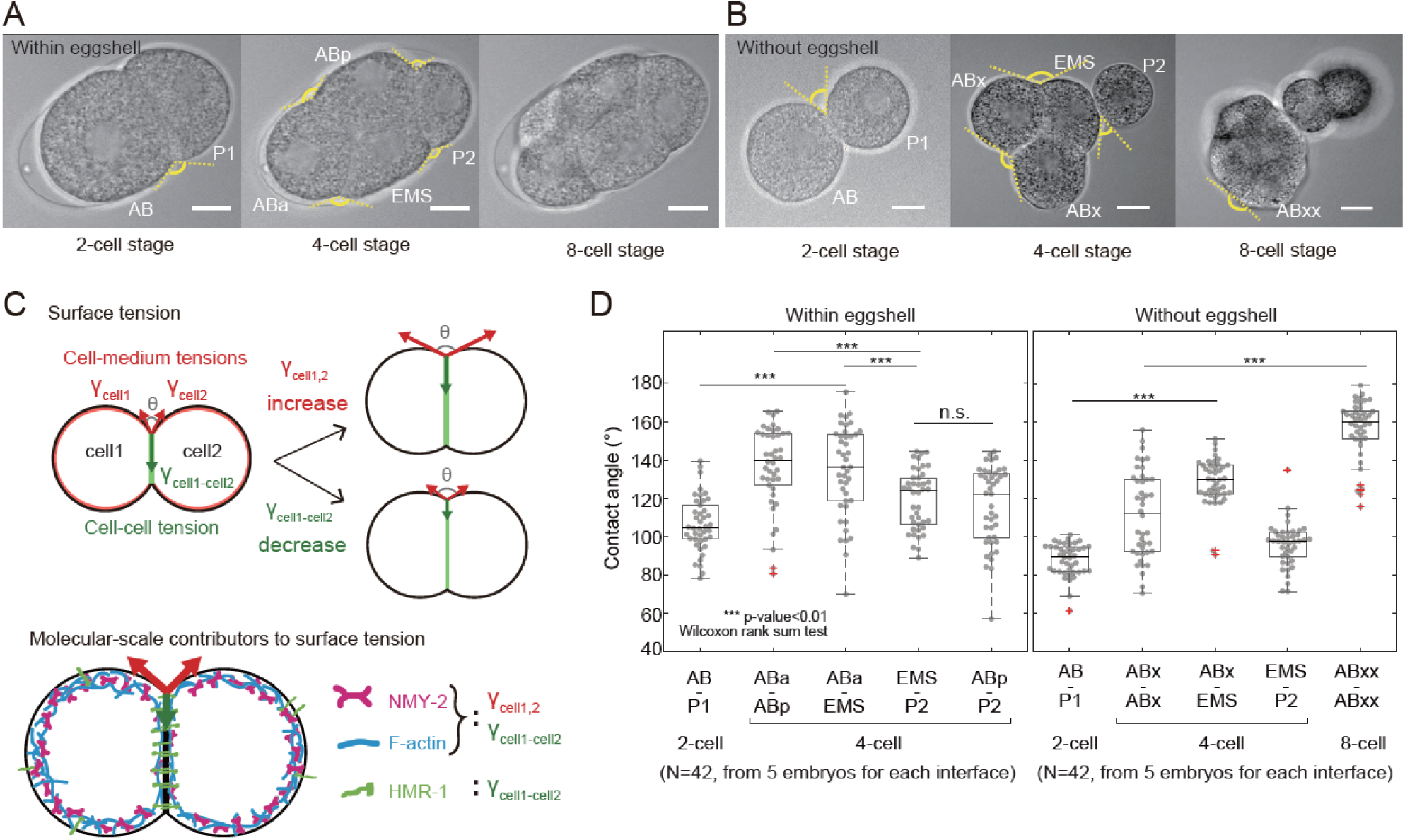
The early *C. elegans* embryo undergoes cell-specific mechanical changes. **A**. Brightfield images of *C. elegans* embryos at the 2-, 4-, and 8-cell stages. Scale bars=10 μm. Dashed yellow lines display the external angle of contact between cells. **B**. Brightfield images of *C. elegans* embryos with the eggshell removed at the 2-, 4- and 8-cell stages. Cell names appear next to each cell in the 2- and 4-cell stage images. Dashed yellow lines display the external angle of contact between cells. Scale bars=10 μm. **C**. Top: The external angle of contact between two cells depends on the balance of cell-medium tension *γ*_cell_ in each cell (red arrows) and the interfacial tension *γ*_cell1-cell2_ at the intercellular contact (green arrow). If the angle of contact increases, this can be due to an increase in *γ*_cell_, a decrease in *γ*_cell1-cell2_, or a combination of both. Bottom: At the molecular-scale, the cell-medium tension *γ*_*cell*_ arises from the action of myosin motor proteins (NMY-2, magenta) on the F-actin scaffold (blue) in the cell cortex. In addition to these proteins, the interfacial tension *γ*_cell1-cell2_ also involves direct and indirect contributions from cell-cell adhesion proteins (E-cadherin or HMR-1, green) and their interactors. **D.** Contact angles measured from brightfield images at different stages of embryonic development in embryos with and without eggshells (n=42, from 5 embryos). Boxplots indicate the 25th and 75th percentiles, the black line within the box indicates the median, and the whiskers extend to the most extreme data points that are not outliers. Outliers are indicated by red crosses. Individual data points are indicated by grey dots. For statistical test, Wilcoxon rank sum test was done.

To gain a first quantitative understanding into the forces shaping the early *C. elegans* embryo, we examined the contact angles between cells. Indeed, the external contact angle *θ* between two cells *i* and *j* is controlled by the balance of tensions in the cell-medium interface of each cell *γ*_*i*_ and *γ*_*j*_ and the tension at their intercellular contact *γ*_*ij*_ (**Fig 1C**). Any change in the angle of contact results from a change in cell mechanics. For example, if the contact angle *θ* between two identical cells increases, this could be because *γ*_*i*_ increases in one or both of the cells, because *γ*_*ij*_ decreases, or because of a combination of both ^34^ (**Fig 1C**). As illustrated schematically, one major challenge in understanding quantitatively multicellular aggregate shape changes is to disentangle the relative contributions of tension at cell-medium interfaces from tension at cell-cell interfaces and to delineate their molecular control by the contractile activity of myosin motors and cadherin adhesion proteins.

During its development, the *C. elegans* embryo is confined in a protective eggshell, which adopts a prolate ellipsoid shape as a first approximation (**Supplemental theory Fig 4A**). Since confinement can force cells against each other, we removed the embryos from their eggshell at the 1- or 2-cell stage to measure angles of contact between cells in brightfield images. This revealed very clear differences in how cells contacted one another (**Fig 1B, S1B**). In excoriated embryos, the positional information that allows to distinguish ABa from ABp is lost and we therefore designate these cells as ABx and their daughters as ABxx. At the 4-cell stage, EMS and the two ABx cells formed a compact aggregate with P_2_ excluded from this and appearing weakly bound to EMS. Later, ABxx cells formed a compact aggregate with MS, while E, P_3_ and C were weakly attached to this group. From a morphological standpoint, cells located at the anterior of the embryo ABx(2)-EMS and ABxx(4)-MS appeared to form increasingly compact clusters of cells, similar to what is observed in 8-cell mouse embryos ^30^. These qualitative relationships were also reflected in the angles of contact between cells. The angle of contact between cells of the P and the AB lineages increased between the 2- and 4-cell stage (**Fig 1D**). Similarly, the angle of contact between ABxx cells was larger than between ABx cells (**Fig 1D**). In the eggshell, the angles between P_2_ and its neighbors were smaller than any of the other angles (**Fig 1D**). Overall, the changes in contact angle and embryo morphology indicated that cells within the embryos possess clear mechanical differences and that their mechanics evolve dynamically even during the earliest stages of morphogenesis.

### Morphogenesis at the 4-cell stage in excoriated embryos is captured by a heterogeneous foam model of the embryo

To advance beyond the previous geometrical and static analysis, we hypothesized that the shape and arrangement of cells outside of the eggshell are primarily controlled by effective surface tensions at each interface in the 2 and 4-cell stage embryos, modeled therefore as heterogeneous foam-like structures. We implemented a tension inference approach (**Fig 2A)**, allowing us to deduce the relative tension values at all interfaces using only measurements of contact angles ^35–38^. We decided not to employ a force inference approach previously applied on *C. elegans* as it solves pressure and tension balance in a combined manner ^39^. This leads to inevitable errors in Laplace pressure because the confinement of the embryo in the eggshell corrupts inferred tension values ^40^. To experimentally image cell geometry, we used a worm line that expresses a GFP-tagged membrane marker (the PH domain of PLCδ) and an mCherry-tagged histone marker to allow determination of the cell cycle stage (**Fig 2A left**, **Fig 2G top**, **Video S1**). From the membrane signal at midplane we could segment the cell outlines (**STAR Methods and Supplemental theory**), and extract the cell contour as non-manifold polygonal lines using a custom-written algorithm, called Delaunay-watershed 2D ^40^; this allowed to measure contact angles at each time-point in each embryo (**Fig 2A, middle**). At junction lines joining three cell interfaces (later called trijunctions), the three surface tensions must balance to ensure mechanical equilibrium. In a 2D midplane slice of the embryo, these trijunctions appear as points where contact angles can be easily measured (**Fig 2A, middle)**. Instead of using the classical Young-Dupré tension balance, which relates contact tensions to the cosines and sines of the contact angles, we derived an equivalent mathematical expression that directly expresses each tension as a function of the cotangents of the contact angles, multiplied by a common factor (**Supplemental Theory Fig 3**). This expression clearly demonstrates that contact angles alone can inform us about the relative values of tensions at cell interfaces, as the common factor cancels out in tension ratios. Knowing the angles at each trijunction within the embryo provides a complete set of linear equations relating all relative tensions. By adding a constraint to set a reference tension at one interface, this linear system becomes overdetermined and can be solved in a least-squares sense using a Moore-Penrose pseudo-inverse (**Supplemental theory**). This allows inference of all surface tensions relative to a reference tension within the embryo, that we chose as the cell-medium tension of P_1_ for 2-cell stage embryos and of P_2_ for 4-cell stage embryos.

**Figure 2:**
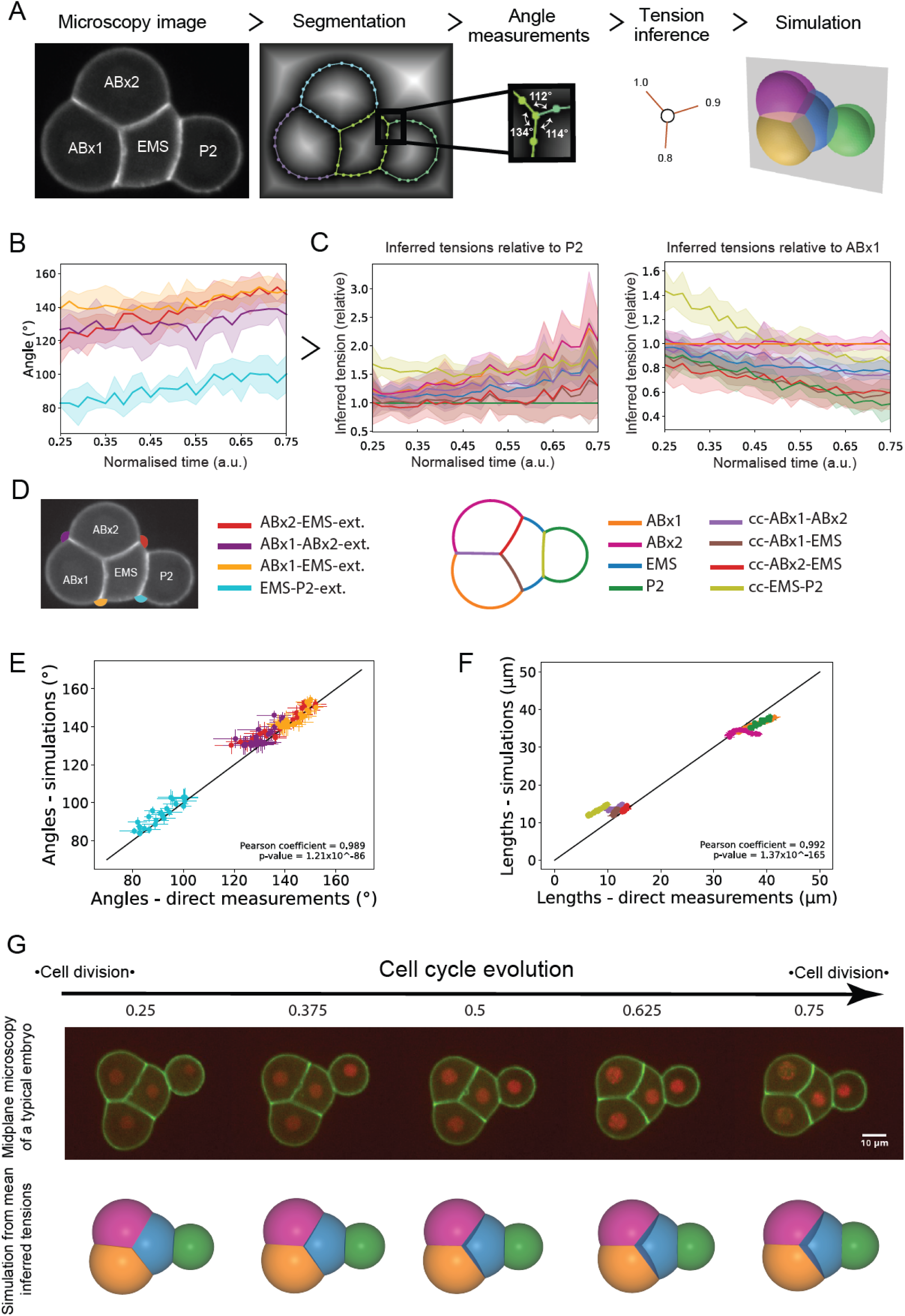
Tension inference allows prediction of spatiotemporal shape changes in the embryo. **A.** Tension inference pipeline illustrated on an embryo outside of the eggshell. Microscopy images of the cell membrane (first column) are segmented and the cell outlines are represented by non-manifold polygonal lines (second column). At each contact point where three cell surfaces meet, the angles between all surfaces are determined (third column). The relative tensions in each surface are then inferred based on the angles of contact (fourth column). 3D heterogeneous foam simulations predict cell arrangement (fifth column) based on the relative tensions inferred at each surface. **B.** Temporal evolution of contact at tricellular junctions. The solid line shows the mean and the shaded area represents the standard deviation (n=5 embryos). Time is normalised to the cell cycle and the color codes relating to external angles of contact are indicated in the panel D below. Internal angles are not shown here for the sake of readability. **C.** Temporal evolution of inferred surface tensions relative to P_2_ (left) and relative to ABx1 (right) for embryos outside of the eggshell. The differences between the two panels illustrates the influence of the choice of temporal reference when reporting relative tension values. The solid line shows the mean and the shaded area represents the standard deviation (n=5 embryos). The color codes relating to cell interfaces are indicated in the panel D below. **D**. Left: Color code for contact angles used throughout the manuscript. Right: Color code used for interfaces throughout the manuscript. **E.** Plot of the contact angles obtained from simulations as a function of experimentally measured angles. Each data point represents one time point with standard deviation and is averaged over 5 embryos. The correlation is measured through a Pearson coefficient ρ=0.989. The black line shows the line of slope 1. **F.** Plot of the simulated lengths as a function of experimentally measured lengths. Each data point represents one time point with standard deviation and is averaged over 5 embryos. The correlation is measured through a Pearson coefficient ρ=0.992. The black line shows the line of slope 1. **G.** Temporal evolution of inferred surface tensions allows prediction of the cell arrangement in embryos. First row: microscopy time series of a developing embryo outside of the eggshell. The membrane is visualized with GFP-PH-PLCδ (green) and the histones are visualized with mCherry::his-58 (red). Scale bar=10µm. Second row: temporal evolution of the mean 3D embryo shape predicted by simulation (n=5 embryos). ABx1 appears in yellow, ABx2 in pink, EMS in blue, and P_2_ in green.

**Figure 3:**
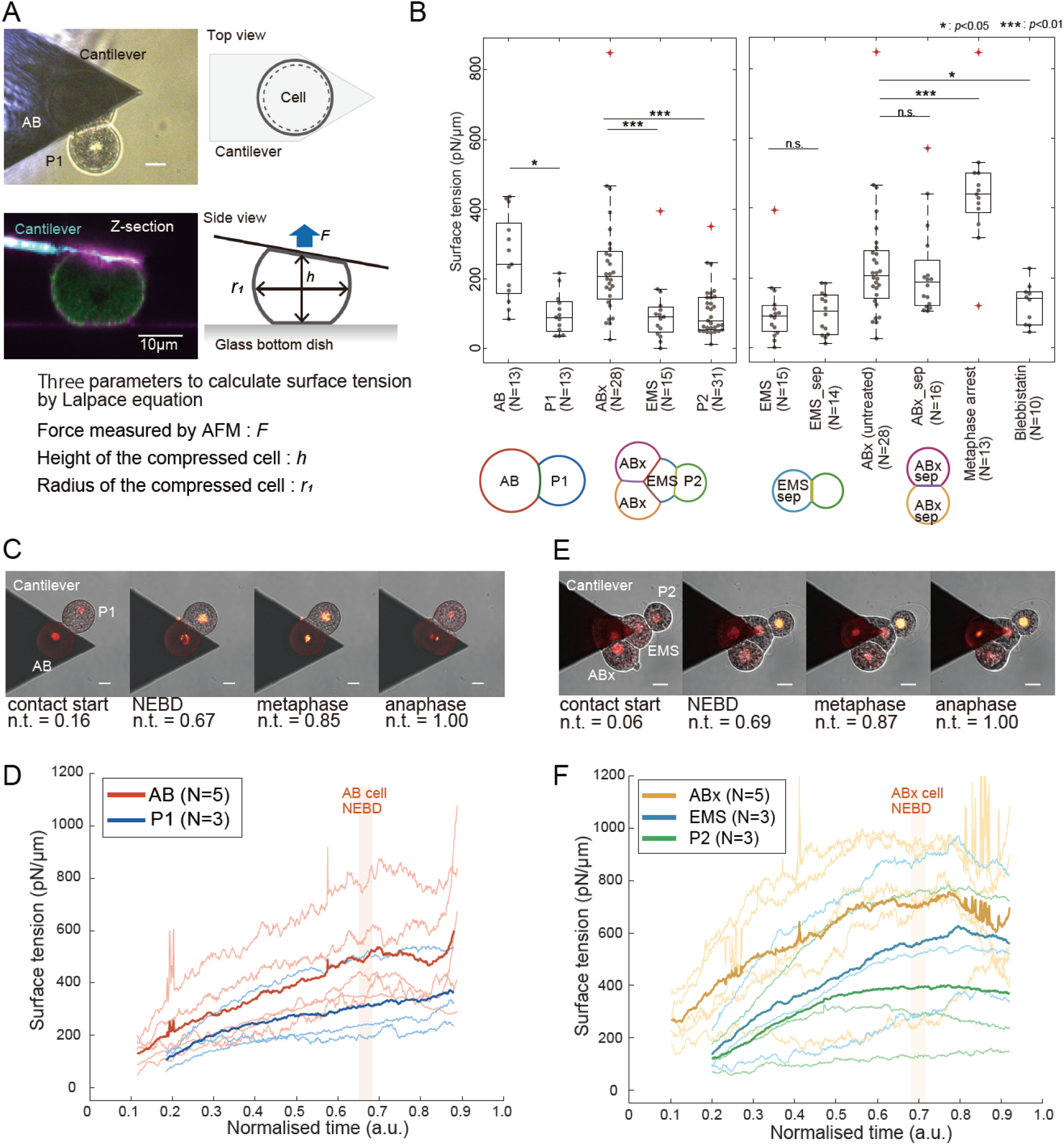
AB and P lineage cells are mechanically different in 2- and 4-cell stage *C. elegans* embryos. **A.** *C. elegans* embryos during AFM measurements of surface tension. Top left: brightfield image of an AFM cantilever in contact with the AB cell in a 2-cell stage embryo. Scale bar= 10μm. Top right: schematic diagram of the measurement. Bottom left: profile view of the AFM cantilever in contact with the AB cell of a 2-cell stage embryo. The cell membrane is labelled with CellMask (magenta) and the embryo expresses NMY-2::GFP (green). The AFM cantilever is visualized by reflection (cyan). Bottom right: diagram indicating the force exerted by the embryo on the AFM cantilever (blue arrow) and the cell dimensions (black double arrow). **B.** Single time point measurements of cortical tension in cells of the *C. elegans* embryo at the 2- and 4-cell stages. Boxplots indicate the 25th and 75th percentiles, the thick black line indicates the median, and the whiskers extend to the most extreme data points that are not outliers. Outliers are indicated by red crosses. Individual data points are indicated by grey dots. Left: Cortical tension in the cells of the embryo at the 2- and 4-cell stages. A diagram of the cellular arrangement in those stages is presented below the graph. Middle: Comparison of the cortical tension in cells from intact embryos and from embryos in which AB and P_1_ were separated. After cell division, tension is measured in the ABx cell (ABx-sep) and in the EMS cell (EMS-sep). Right: Cortical tensions in ABx cells in untreated embryos and in embryos treated with blebbistatin to inhibit myosin contractility or arrested in metaphase by proteasome inhibition with c-lactocystin-ß-lactone. The number of cells from different embryos in each condition is indicated on the x-axis. Different conditions are compared with a Wilcoxon rank-sum test. **C.** Time series of a 2-cell embryo during tension measurement. The cantilever is visible as a triangular dark shadow and is in contact with the AB cell. The GFP:: H2B signal is overlaid on the brightfield microscopy images in red, allowing determination of the phase of the cell cycle. **D.** Temporal evolution of cell-medium tension in AB (average of n=5 embryos) and P_1_ cells (n=3 embryos). The thick line indicates the mean and the thin lines show each individual measurement. Time was normalised to the cell cycle. The timing of NEBD in AB is indicated. (**E-F)** Same as (C-D) but for a 4-cell embryo. Evolution in ABx (average of n=5 embryos), EMS (n=3 embryos) and P_2_ cell (n=3 embryos). The timing of NEBD in ABx is indicated.

To account for differences in development duration across embryos, we normalised time choosing the moment of closure of the cytokinetic ring of the mother cell as time 0 and the beginning of furrowing in the next cell cycle as time 1 (**Fig S1A,B**). From cell outlines, we extracted and plotted the time evolution of contact angles (**Fig 2B**). At the 4-cell stage, all external angles of contact grew by ∼10-20°, although their temporal dynamics were somewhat different. These data confirmed that dynamic changes in mechanics took place in all cells of the embryo.

Next, we inferred relative surface tension maps within the embryos without eggshells based on contact angle measurements. To illustrate the necessity of a mechanical reference for accurately inferring the temporal evolution of surface tensions, we compared the temporal evolution of inferred tensions by arbitrarily keeping either the cell-medium tension of P2 or ABx1 constant over time (**Fig 2C**). In the former case, all tensions increase over time, while in the latter, all tensions decrease except for ABx2, which remains roughly constant as expected due to its indistinguishability from ABx1. This comparison demonstrates that the tension inference method alone, static and relative by nature, is not sufficient to infer the temporal dynamics of surface tensions, and a reference is necessary to calibrate tension values in time from the evolution of their ratios. Nevertheless, relative tension values remain sufficient to fully characterize cell shape and arrangement at any timepoint, which we demonstrate by re-simulating cell shape and arrangement from inferred tensions (**Fig 2A**) using a previously developed 3D heterogeneous foam computational model ^16,30^. The predicted mean embryo shape not only qualitatively resembles experimentally observed shapes (**Fig 2G, Video S2**), but the mean geometry of simulated cells agrees quantitatively very well with experimental ones, as measured by the comparison of interface lengths in the embryo midplane (**Fig 2F**) and contact angles (**Fig 2E**). Similar agreement between prediction and experiment was also obtained for the 2-cell stage (**Fig S2A-E, Video S2)**. The close match between experimental measurements and simulations indicates that a static heterogeneous foam model can effectively describe the average shape evolution of *C. elegans* embryos in the 2- and 4-cell stages. This result validates our use of the tension inference method to characterize the evolution of embryo shape in each stage.

### Cell-medium tension differs across cell lineages

To go beyond relative information on tensions across interfaces, we decided to directly measure the cell-medium tension *γ*_*i*_ in each individual cell in 2- and 4-cell stage embryos using Atomic Force Microscopy (AFM) (**see STAR Methods and Supplemental Theory, Fig 3, Fig S3A-K**). As these measurements require direct contact between a microfabricated cantilever and the cell, we excoriated the embryos, manually removing the embryonic eggshell. AFM-based tension measurements are usually carried out on isolated cells but, as a first step, we assumed that the presence of a cell-cell contact on part of the cell periphery did not significantly affect estimation of cell-medium tension. In our experiments, we brought a tipless AFM cantilever into contact with a cell and waited 30s for the measured force to relax to a plateau (**Fig S3A**) ^41^. This allowed passive stresses originating from deformation of the cortical F-actin network to be dissipated by turnover and ensured that we measured only steady-state tension arising from active mechanisms, such as myosin contractility ^41^. The force at the plateau together with the geometry of the cell allow to estimate the cell-medium tension by explicitly solving the Laplace equation along the lateral cell profile, assuming negligible adhesion of the cell with the substrate ^42^ (**Methods and Supplemental theory**).

The mean cortical tension in *C. elegans* cells ranged between ∼100-300 pN/μm (**Fig 3B**), comparable in magnitude to that reported in interphase in mammalian cultured and embryonic cells ^8,43^ as well as in invertebrate embryos ^18^. Interestingly, clear mechanical differences were evident between cell lineages. Cells of the AB lineage (AB and ABx) had tensions ∼2.5-fold larger than the P lineage cells (P_1_, P_2_) (**Fig 3B, left**). As previously observed in mammalian cells, cortical tensions presented a large variation around the mean, perhaps due to embryo-to-embryo variability or differences in cell cycle progression. To determine if the presence of intercellular contacts affects cell-medium cortical tension estimation, we separated P_1_ from AB at the 2-cell stage, waited for each of these cells to divide, and measured the cortical tension in ABx_sep_ and EMS_sep_. We found no significant difference in cortical tension between ABx, which has 2 contacts, and ABx_sep_, which has only one, or between EMS, which has 3 contacts, and EMS_sep_, that only has one (**Fig 3B, right**). This suggests that the presence of contacts has a less significant effect on cell-medium cortical tension estimation than other sources of regulation or variability.

One potential reason for the observed mechanical differences between the AB and P lineages is the asymmetric partitioning of actomyosin between AB and P_1_ that takes place during the first cell division ^9,10^. Indeed, in most model systems, the submembranous actomyosin cortex is a key determinant of cell surface tension ^30,44^. As AB inherits most of the actomyosin cortex of P_0_, we would expect it to have a larger surface tension than P_1_. When we imaged the actomyosin content in 4-cell stage embryos, we could observe clear differences in myosin intensity with ABa and ABp being brighter than EMS and P_2_ (**Fig S1H, I**), whereas differences in F-actin staining were more subtle (**Fig S1H, J**). The same trend was confirmed between AB and P_1_ for 2-cell stage embryos (**Fig S1E-G**). This suggested that, as in other model systems, myosin contractile activity may play a key role in setting cellular mechanical properties. Consistent with this, inhibition of myosin contractility with blebbistatin decreased cortical tension in ABx cells and, conversely, metaphase arrested ABx cells, in which myosin activity is upregulated, displayed increased cortical tension (**Fig 3B, right**). Together, these experiments reveal clear differences in cell-medium cortical tension between cells in the embryo that may result from asymmetric partitioning of myosin over successive divisions.

### Cortical tension evolves dynamically during the cell cycle

In our single time-point measurements (**Fig 3B**), we observed a wide range of cortical tensions in each cell type. Given the clear increase in tension during metaphase, we hypothesized that this variability may arise from the combination of the dynamic evolution of cell-medium cortical tension *γ*_*i*_ throughout the cell cycle together with our static measurements occurring at different stages of this cycle. To determine temporal changes in cell-medium cortical tension, we compressed cells with an AFM cantilever and maintained contact during the whole cell cycle while simultaneously imaging GFP::H2B localization to follow cell cycle progression (**Fig 3C, E, Video S3**). We monitored the restoring force exerted on the AFM cantilever as this reflects changes in *γ*_*i*_. In all cell types, we observed an increase in restoring force with magnitudes far larger than the measurement sensitivity of AFM in our experimental conditions (>3nN increase vs ∼0.1nN sensitivity, **Fig S3B-F**), signifying that we could accurately quantify the evolution of *γ*_*i*_ during interphase (**Fig 3D, F and Fig S3G-K**). At the 2-cell stage, *γ*_*AB*_ increased ∼2-fold at an approximately constant rate over the course of interphase, reaching a maximum at Nuclear Envelope Breakdown (NEBD) (red line, 5/5 cells, **Fig 3D, Fig S3G**). In anaphase, the force applied on the cantilever increased dramatically because the AB cell often divided perpendicular to the plane of the coverslip (**Fig S3B**). Note that it was not possible to determine the change in tension in anaphase because analytical models assume a spherical geometry (**Supplemental Theory**). *γ*_*P*1_ was always lower than *γ*_*AB*_ but it also increased ∼2-fold (blue line, 3/3 cells, **Fig 3D, Fig S3H**). No large change in force was observed at anaphase in P_1_ because it always divided in the plane (**Fig S3C**). At the 4-cell stage, the ABx cells displayed a ∼1.5-fold increase in tension (orange line, 5/5 cells, **Fig 3F, FigS3J**). *γ*_*EMS*_ was initially ∼2-fold smaller than *γ*_*ABx*_ at the beginning of its cell cycle but increased over 3-fold, reaching a value comparable to *γ*_*ABx*_by NEBD (light blue line, 3/3 cells, **Fig 3F, Fig S3K**). This dramatic change in cell-medium tension may underlie the compaction of ABx(2)-EMS observed at the end of the 4-cell stage (**Fig 1A,B**). In contrast, *γ*_*P*2_, which was initially similar to *γ*_*EMS*_, only increased by ∼2-fold during its cell cycle (green line, 4/4 cells, **Fig 3F, Fig S3L**). Overall, cell-medium tension increased in all cells during interphase at both the 2- and 4-cell stages (**Fig S3I, M**).

Overall, our AFM measurements show that all *C. elegans* blastomeres increase their cell-medium tension during their cell cycle at 2 and 4-cell stage. The larger cell-medium tension in ABx coupled with the large increase in *γ*_*EMS*_ in the anterior of the embryo may drive the compaction of ABx(2)-EMS. The steady increase in surface tension observed during the cell cycle (**Fig S3G-M**) contrasts with the constant surface tension reported in mammalian cells in metaphase over a similar duration ^41,45^ or with the pulsatile contractility observed in compacting mouse embryos ^30^.

### Cortical enrichment in myosin correlates with inferred and measured cell-medium tensions

While the measurement of one cell-medium tension may allow calibration of tensions over time in embryos outside of their eggshells, characterizing cell mechanics within the eggshell is not possible with AFM. Additionally, whether cortical tension is identical inside and outside of the eggshell remains unclear. This question is especially relevant for ABp, which interacts with P_2_ through Delta-Notch signaling due to a contact arising as a consequence of shell confinement that drives ABp towards a different fate than ABa ^14,15^. Therefore, as an alternative, we sought to identify molecular markers that could be used to temporally calibrate surface tensions within the eggshell. As the AB lineage had the highest cortical tension in our AFM measurements (**Fig 3**) and inherits most of the actomyosin ^9^ (**Fig S1)**, we hypothesized that cortical myosin fluorescence may be a good proxy for cortical tension, as in other systems such as mouse, *Drosophila* or zebrafish. To test this, we acquired time-lapses of embryos expressing a GFP fusion to the endogenous NMY-2 gene, which encodes non-muscle myosin heavy chain, and segmented the data to obtain the temporal evolution of myosin enrichment at the cortex of each cell and at each cell-cell contact.

At 2-cell stage when the eggshell was removed, myosin fluorescence intensity in the cortex of AB was ∼2-fold higher than in P_1_ (**Fig 3A**, **Fig S1F, Fig S4C**), consistent with the ratio of their cortical tensions measured by AFM (**Fig 3B, D**). At the 4-cell stage, the ABx cells had the highest cortical myosin fluorescence, followed by EMS and finally P_2_ (**Fig 4D**, **Fig S4D**), similar to the ordering of cortical tensions measured by AFM (**Fig 3B, F**). Similar trends could also be observed for embryos within their eggshell at the 2- and 4-cell stages (**Fig S4A, B, Video S4**). Overall, these data supported the hypothesis that cell-medium myosin cortical enrichment reflects cell-medium cortical tension. As NMY-2 fluorescence is only indicative of total myosin, we immunostained embryos against the phosphorylated, active myosin, which generates tension (p-MLC, **Fig 4B**). This revealed that ABx cells had the highest cortical signal, followed by EMS, and P_2_, similar to the ordering of cortical tensions measured by AFM. In addition, the NMY-2 and p-MLC signals showed a high degree of correlation in all of the embryonic surfaces (**Fig 4C**). When we quantified the temporal evolution of cortical myosin, we found that cortical myosin intensity at cell-medium interfaces increased over the cell cycle (**Fig 4D, Fig S4A-D, Video S4**) similar to the temporal evolution of cortical tensions measured by AFM (**Fig 3F**). Therefore, total myosin appeared as a reasonable proxy for active myosin and differences in myosin intensity across cells appeared well correlated with the cell-medium tensions measured by AFM (black line, **Fig S4E**). However, when we attempted to use this correlation directly in simulations, we could not predict cell arrangement in embryos. We reasoned that this is because AFM measurements only measure tension evolution in one cell in each embryo and because there is a large variability in myosin expression (**Fig S4A-D**) and cell-medium tension across embryos (**Fig 3D, F**). As a result, average cell-medium tensions determined from separate embryos may not satisfy overall mechanical equilibrium at the embryo scale. These observations suggested that robust regulation of *relative* surface tensions may be key for a robust control of cell arrangement in early embryos.

**Figure 4:**
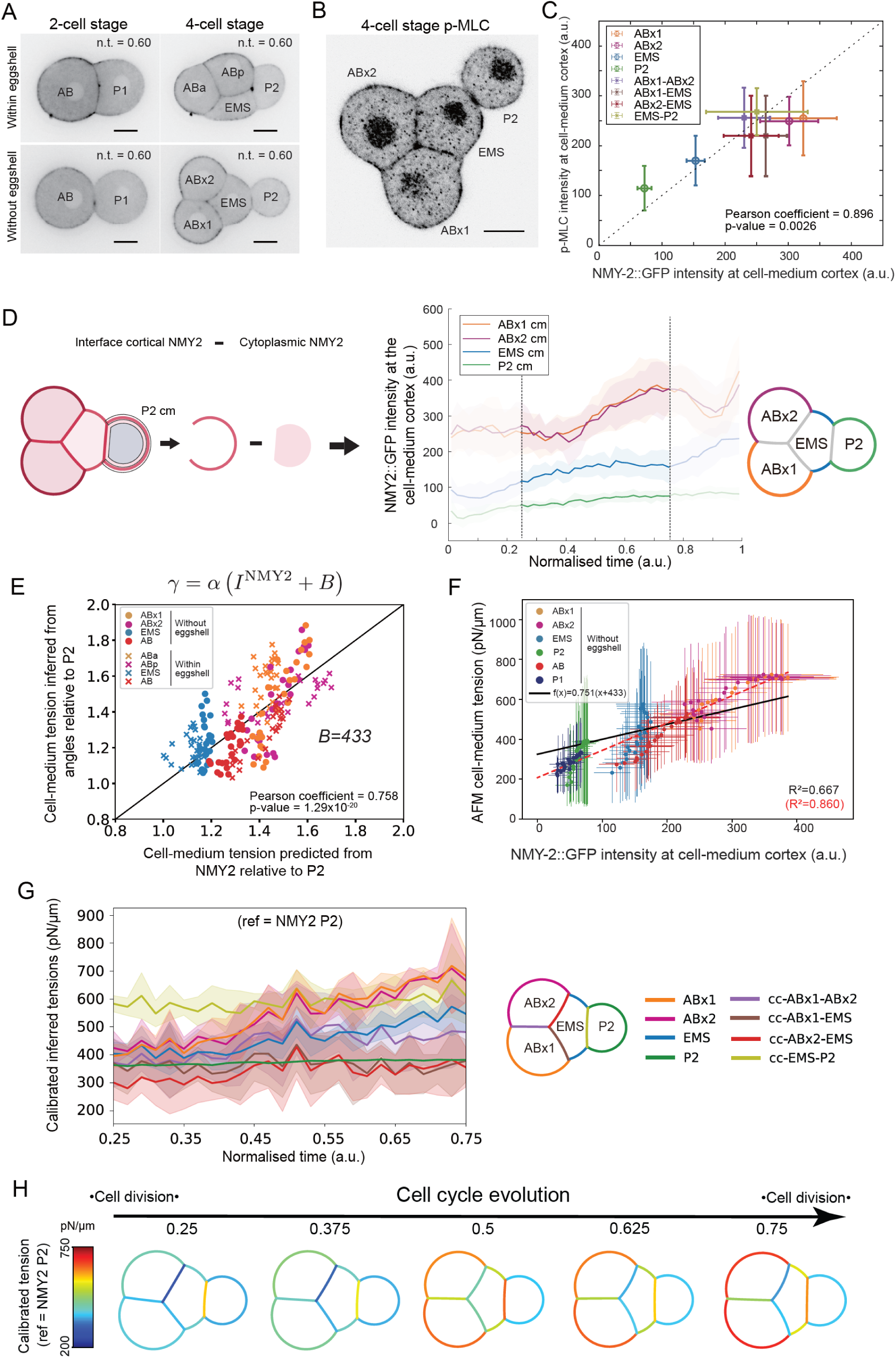
Myosin fluorescence intensity correlates with inferred and measured tensions. **A**. NMY-2::GFP localization in embryos at the 2- and 4-cell stages with and without eggshell at n.t.=0.60. Scale bar= 10μm. The identity of each cell is indicated on the images. Identical intensity histograms were chosen for all images. **B**. Immunostaining against phosphorylated myosin light chain (p-MLC) at the 4-cell stage in an embryo outside of the eggshell. The identity of each cell is indicated on the image. Scale bar=10μm. **C**. Phosphorylated-myosin light chain (p-MLC) fluorescence intensity as a function of NMY-2::GFP fluorescence intensity in all of the surfaces of 4-cell stage embryos. Data averaged over 5 embryos. The whiskers indicate the standard deviations. The correlation is measured by a Pearson coefficient ρ=0.90 and its significance is measured by a p-value=0.003. The dotted diagonal line indicates perfect correlation. **D**. (Left) Sketch of the methodology employed to quantify myosin intensity at the cell-medium cortex with NMY-2::GFP at each cell-medium interface. The average cytoplasmic intensity was subtracted from the average intensity in the cell interfaces to compute the average intensity in the cell-medium cortex. (Middle) Myosin intensity as function of time for all cell-medium cortices in embryos outside of the eggshell. The solid line represents the mean over 5 embryos and the shaded area is the standard deviation. (Right) Sketch of the surfaces where measurements of NMY2::GFP were taken with each surface colour coded. **(E-F)** Data points were taken between n.t.=0.25 and n.t.=0.75. **E**. Relative cell-medium tension inferred from angles as a function of the relative tension predicted from cell-medium myosin fluorescence intensity, where we hypothesized a global relationship 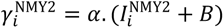. For both axes, the values were normalised to the respective value for P_2_. Each data point represents one time point and is averaged over 5 embryos. The value of *B* = 433 optimizes the linear correlation, measured by a Pearson correlation coefficient ρ=0.758. The solid black line indicates perfect linear correlation. **F**. Cortical tension measured by AFM as a function of the cell– medium myosin fluorescence intensity. Each data point corresponds to a single time point and represents the average over 5 embryos, with standard deviation indicated. The solid black line shows the linear regression under the constraint β = α·B on the intercept, where B = 433 ensures consistency between angle-inferred and myosin-derived tensions (as shown in Fig. 4E). This yields a slope of α = 0.751 pN/(μm·ua) and an intercept β = 325 pN/μm with a coefficient of determination R^2^=0.667. For comparison, the unconstrained linear regression yields R^2^ =0.86 (red dashed line, also presented in Fig. S4E). **G**. Temporal evolution of inferred surface tensions calibrated in time and absolute value using the myosin fluorescence intensity in P_2_ and the affine parameters from F. The solid line shows the average over 5 embryos and the shaded area represents the standard deviation. Time is normalised to the cell cycle. Right: Schematic diagram of the 4-cell stage embryo with each surface color coded. **H**. Temporal changes in absolute mean surface tensions obtained from myosin-informed inference using affine parameters from F. The tension in each surface is color coded with blue representing low tension and red high tension.

To overcome this challenge, we implemented a two-step hierarchical procedure to predict absolute cell-medium tension from cortical myosin intensity. First, in each cell *i*, we searched for a correlation between relative myosin intensities 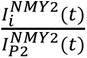 and relative tensions determined from inference 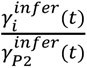 because this analysis inherently satisfies mechanical equilibrium in the whole embryo. Second, we used AFM tension measurements to set an absolute scale for the previous relationship. Our ultimate goal is to predict absolute cell-medium tension from myosin intensity using the relationship: *γ*^*NMY*2^ = *⍺*(*I*^*NMY*2^ + *B*). In this relationship, *⍺* reflects how much each unit of cortical myosin fluorescence contributes to cortical tension and *B* is a myosin-independent contribution to tension, which may reflect the membrane and/or spectrin tension ^46^. To determine *B*, we first performed a linear regression between cell-medium tensions inferred from angles relative to the cell-medium tension in P_2_, and tension values predicted from NMY-2 intensity from the affine relation, relative to predicted value for P_2_. The linear regression of the set of points 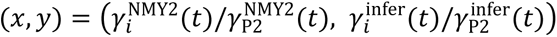 predicts a ratio *B* = 433 that gives the best correlation between relative inferred tensions and relative predicted tensions from NMY-2 intensity both inside and outside of the eggshell (**Fig 4E**). This suggests that myosin intensity serves as a reliable proxy for predicting relative cell-medium tensions both inside and outside the eggshell. It also reveals a constant contribution to cell-medium tension that is independent of NMY-2–regulated contractility, potentially arising from membrane or spectrin tension. This myosin-informed inference approach allowed us to calibrate in time the relative tensions inferred from geometry, but the knowledge of *B* was not sufficient to give them a proper absolute scale.

To provide an approximate absolute scale for our inferred tensions, we performed a linear regression between AFM-measured cell-medium tensions 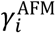 and myosin-derived cell-medium tension intensity 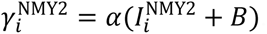. The slope *⍺* was treated as a free parameter while the intercept was constrained as *β* = *⍺* ∗ *B* with *B* = 433 optimized previously in (**Fig 4E**). This yielded *⍺* =0.751 pN/(μm.au) and *β* =325 pN/μm, (**Fig 4F**), enabling absolute calibration of inferred tensions in pN/μm. This constrained linear regression (black line, **Fig 4F**) is not as good as the free regression (red dashed line, **Fig 4F**) because we prioritised the correlation of NMY-2 intensity with inferred tensions to ensure global balance of tensions. However, this approach captured the ordering of tensions in the different cell types and their temporal evolution (**Fig S3N-P**). For some cells (P_1_, AB, and ABx), the cell-medium tension relationship with cortical myosin fluorescence follows the global trendline (see slopes on **Fig S4E, Fig S3P**). Other cells (EMS and P_2_) appeared to possess a steeper slope. As cortical tension depends both on cortical myosin enrichment as well as organization of F-actin and crosslinkers in the cortex^43,44^, these differences in slope may reflect differences in F-actin organisation in the cortex of EMS and P_2_ or a lesser degree of correlation between phosphorylated and total myosin for this lineage. Nevertheless, the relationship allowed to capture a gross estimate of the absolute cell-medium tension values based solely on NMY-2 intensity at the cortex.

With this approach, we could draw a first panorama of the spatiotemporal changes of all absolute surface tensions shaping the 2-cell (**Fig S2F**) and 4-cell embryo outside of the eggshell (**Fig 4G-H)**. At the 4-cell stage, all of the cell-medium cortical tensions with the embryo increased with time but the cortical tensions in ABx(2) and EMS increased the most. Cell-cell surface tensions changed less than cell-medium cortical tensions but each intercellular contact had a very specific magnitude of interfacial tension with *γ*_EMS-P2_ the largest, followed by *γ*_ABx-ABx_, and finally *γ*_EMS-ABx_ (**Fig 4G-H**). Compaction took place in the anterior part of the embryo because cortical tensions were larger than intercellular contact tensions (n.t.=0.4-0.77, **Fig 4G-H**). In the posterior, the cortical tension in P_2_ was always much lower than the tension at its interface with EMS, leading to it bulging out from the EMS-ABx(2) aggregate, especially at early times (**Fig 4G-H**). Thus, the relative magnitude of cell-medium to cell-cell tensions appeared to play a significant role in controlling the characteristic shape of the 4-cell embryo, a phenomenon likely mediated by intercellular adhesions.

To illustrate how spatiotemporal maps can help to understand inhibitor experiments, we treated embryos outside of the eggshell with blebbistatin, an inhibitor of myosin. We chose a moderate concentration (20μM) to achieve partial inhibition of contractility without perturbing processes such as cytokinesis. At the beginning of the 4-cell stage for n.t=0.25, treated and control embryos appeared morphologically similar (**Fig S4F**). Later for n.t=0.6, when compaction of ABx(2)-EMS took place, treated embryos appeared less compacted than control embryos. Consistent with this, significant differences in external angles of contact only appeared later in the 4-cell stage (**Fig S4G**). Although the lack of impact of blebbistatin treatment early in the 4-cell stage may appear surprising, it can be understood from our spatiotemporal surface tension maps (**Fig 4G-H**). Treatment with blebbistatin affects both cell-medium and cell-cell tensions and the external angle of contact is controlled by the ratio of cell-medium tensions *γ_i_* and *γ_j_* and the cell-cell tension *γ_i-j_* (**Fig 1C**): 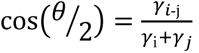.

At the beginning of the 4-cell stage for n.t=0.25, cell-medium and cell-cell tensions are of similar magnitude (**Fig 4H**), suggesting they will be affected similarly by blebbistatin inhibition. As a result, the angle of contact remains unchanged. Later, cell-medium tensions become larger than cell-cell contact tensions because of myosin recruitment (**Fig S4D**). As a consequence, blebbistatin treatment has a larger impact on cell-medium tensions, 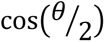 increases, and the angle of contact decreases because cosine is a decreasing function of θ (n.t=0.77, **Fig 4H**).

### Morphogenesis in the eggshell stems from dynamic changes in surface tensions

Next, we applied the same approach to morphogenesis within the eggshell, for which we cannot obtain direct experimental tension measurements. We assumed therefore that the relationship between cortical cell-medium tension and cortical myosin intensity remained similar. Again, we imaged a worm strain expressing GFP-tagged membrane and mCherry-tagged histones, determined relative surface tensions from measurement of the angles of contact between cells, and calibrated their absolute magnitude using our affine relationship (**Fig 5A)**. We extended our foam simulations to predict cell shape and arrangement in the embryo in the presence of a rigid eggshell confinement by implementing a non-penetrating ellipsoidal boundary parametrized from the average shape of the cuticle (**Supplemental theory Fig 4, Fig S3A**). As input, we provided the temporal evolution of mean relative tensions at all interfaces inferred from angles of contact. The predicted embryo morphologies quantitatively followed experimental observations, as illustrated by the comparison of simulated and mean measured interface lengths at the midplane (**Fig 5B**) and the comparison of simulated and mean measured contact angles (**Fig 5C**). Qualitatively, simulations using the mean relative tensions in each surface as input clearly displayed the transition from a loosely linked embryo at the beginning of the 4-cell stage to a more tightly packed one at the end (**Fig 5E, middle row**). Overall, this indicates tension inference is sufficient to predict the morphogenesis of the early embryo confined within its eggshell and validates our foam model hypothesis for wild-type embryos. Inference, simulation and validation was performed for each individual embryo and similar results were also obtained for the 2-cell stage (**Fig S5)**.

**Figure 5:**
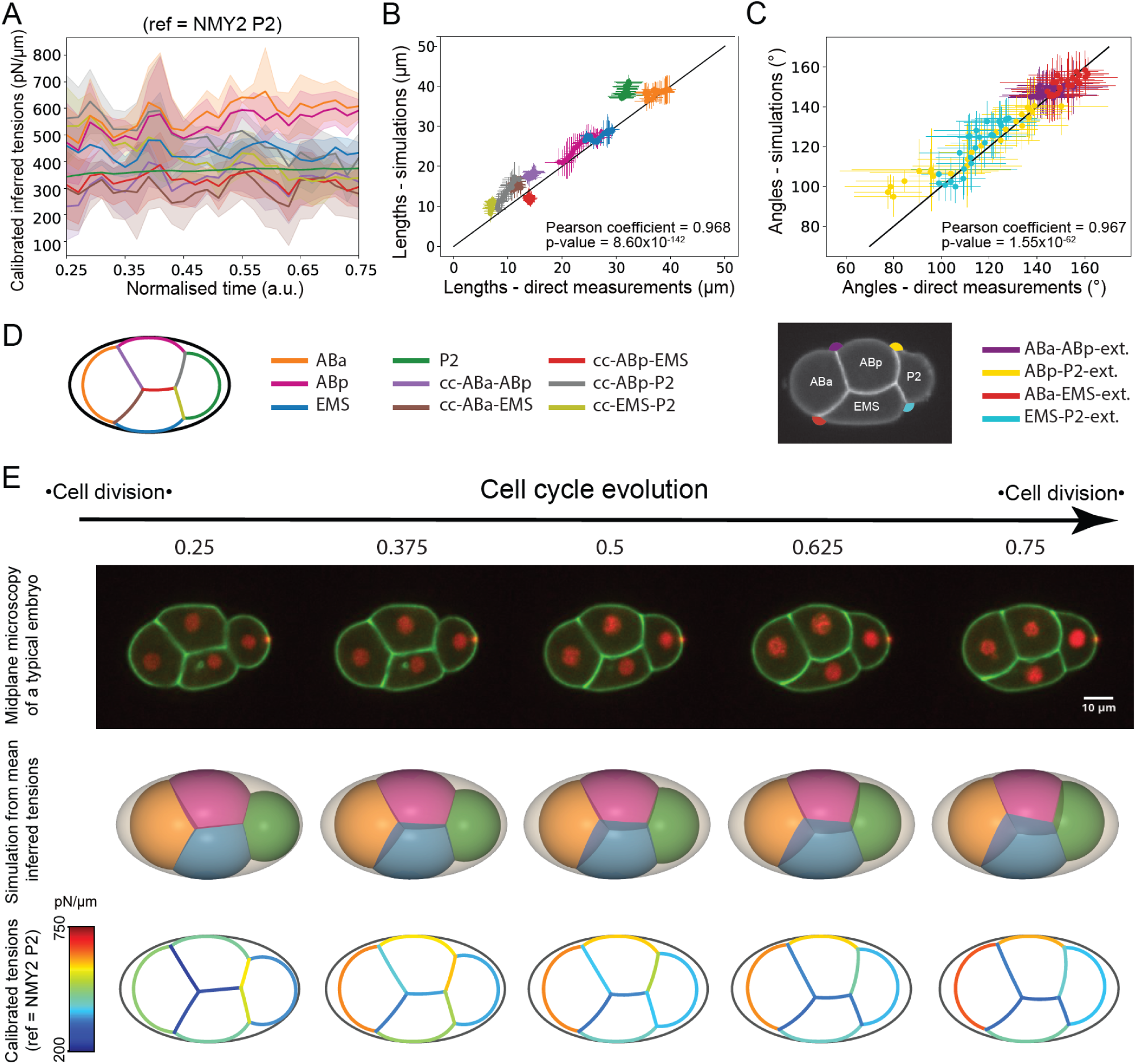
Dynamic regulation of surface tensions underlies morphogenesis in the eggshell. **A.** Temporal evolution of the mean inferred surface tensions calibrated in time and absolute value with the myosin fluorescence intensity in P2 and the affine parameters α=0.751 pN/(μm.ua), β=325 pN/μm. The solid line shows the average over 5 embryos and the shaded area represents the standard deviation. Time is normalised to the cell cycle. Colour code is indicated in D. **B.** Plot of the contact angles obtained from simulations as a function of experimentally measured angles. Each data point represents one time point with standard deviation and is averaged over 5 embryos. The correlation is measured through a Pearson coefficient ρ=0.968. The black line shows the line of slope 1. **C.** Plot of the simulated lengths as a function of experimentally measured lengths. Each data point represents one time point with standard deviation and is averaged over 5 embryos. The correlation is measured through a Pearson coefficient ρ=0.967. The black line shows the line of slope 1. **D**. Right: Color code for contact angles used throughout the manuscript. Left: Color code used for interfaces throughout the manuscript. **E**. Temporal evolution of inferred tensions allows prediction of the cell arrangement in embryos within the eggshell. Top row: microscopy time series of a representative 4-cell embryo developing inside the eggshell. The membrane is tagged with GFP-PH-PLCδ (green) and the histones with mCherry::his-58 (red). Scale bar=10μm. Middle row: temporal evolution of the mean 3D embryo shape predicted by simulation in the eggshell. ABa appears in yellow, ABp in pink, EMS in blue, and P_2_ in green. The eggshell appears in grey (n=5 embryos). Bottom row: mean temporal changes in mean absolute surface tensions obtained from myosin-informed inference using affine parameters α=0.751 pN/(μm.ua), β=325 pN/μm. The tension in each surface is color coded with blue representing low tension and red high tension (n=5 embryos). The eggshell is represented in black.

Then, we combined the evolution of relative tensions with our relationship between cortical cell-medium tension and myosin fluorescence to obtain a spatiotemporal map of absolute surface tensions in the whole embryo (**Fig 5E, bottom row**). This confirmed that compaction of ABa, ABp, and EMS in the anterior of the embryo was driven by a larger increase in cell-medium cortical tensions relative to tension at cell-cell contacts. P_2_ was initially segregated from this group because the intercellular contact tensions *γ*_EMS-P2_ and *γ*_ABp-P2_ were larger than the cortical tension *γ*_P2_. Over time, P_2_ packed more tightly with the rest of the aggregate because intercellular contact tensions decreased, presumably because of the combination of an increase in cadherin at those contacts (**Fig S6B**) and a decrease in NMY-2 (**Fig 6B**). As the decrease in NMY-2 appeared concomitant with the increase in E-cadherin, it may represent an indirect contribution of cadherins to the control of cell-cell surface tension through regulation of myosin contractility ^12,31^.

**Figure 6:**
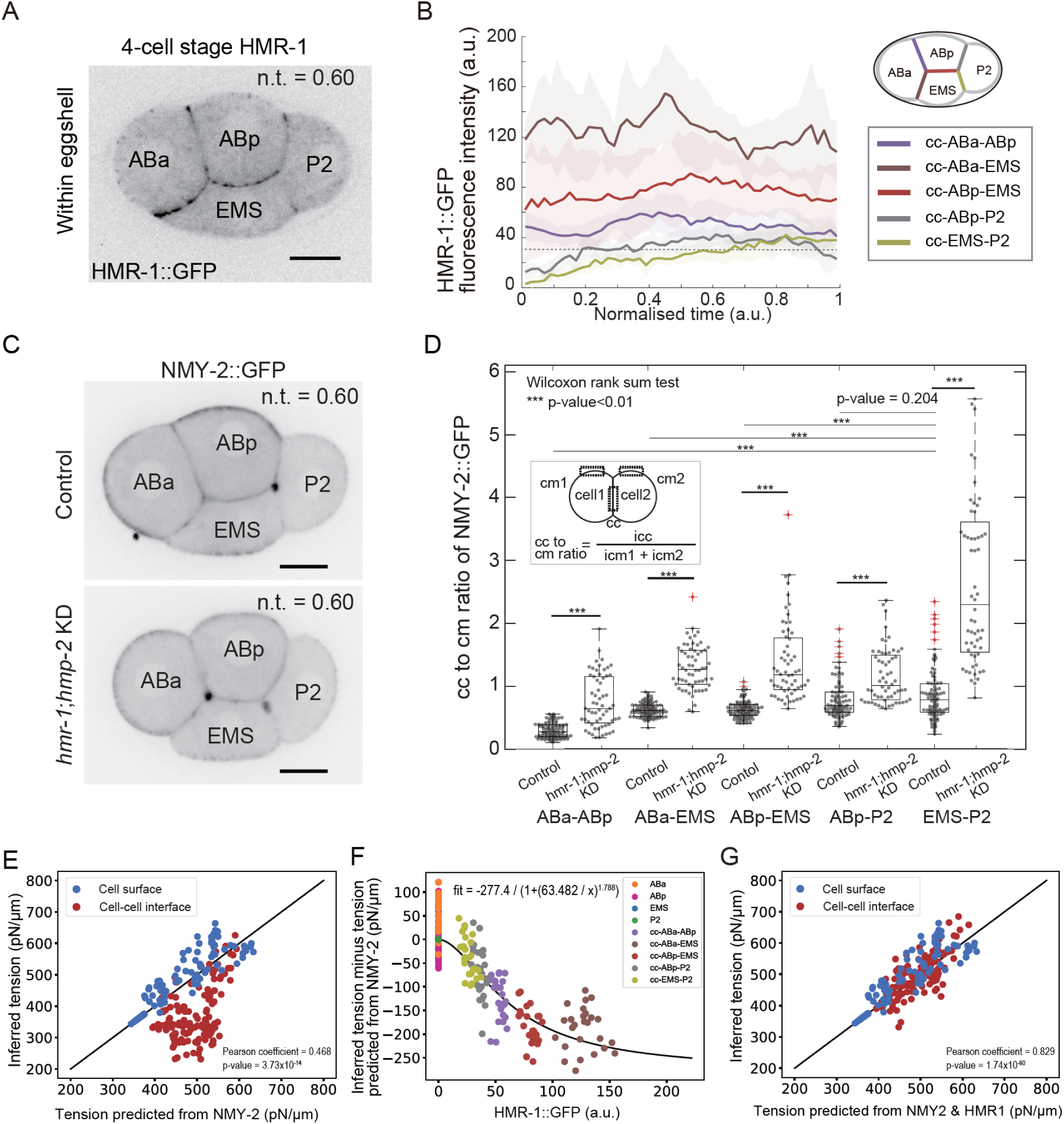
E-cadherin contributes directly and nonlinearly to the tension at intercellular contacts. **A**. E-cadherin-GFP (HMR-1) localization in a representative embryo at the 4-cell stage within an eggshell. Scale bar=10μm. The identity of each cell is indicated on the image. **B**. Temporal evolution of HMR-1 fluorescence intensity in 4-cell stage embryos within the eggshell. The solid line is the average over 5 embryos and the shaded region depicts the standard deviation. All intensities are given in a.u. The dashed line indicates the intensity below which the enrichment at intercellular junctions is visually indistinguishable from background. **C**. NMY-2::GFP localization in a representative embryo at the 4-cell stage within an eggshell in control (upper) and *hmr-1;hmp-2* KD condition (below). Scale bar=10μm. The identity of each cell is indicated on the image. **D**. Comparison between NMY-2::GFP fluorescence intensity at intercellular contacts and at the cell-medium cortex in each pair of cells in contact in control and *hmr-1;hmp-2* KD condition. Top left: sketch of the measurement. The ratio is computed as the intensity I_cc_ in the cell-cell contact divided by the sum of the intensities I_cm1_ and I_cm2_ in the cortices. Temporal averages of the ratio of intercellular to cortical NMY-2::GFP intensities for cell-cell contacts in the 4-cell stage embryo taken between n.t.=0.25 (ABa-ABp, ABa-EMS, ABp-EMS cell-cell interfaces) or n.t.=0.4 (ABp-P2, EMS-P2 cell-cell interfaces) and n.t.=0.75. Data from 5 different embryos and 18 or 13 time points per embryo. Data was taken between n.t.=0.4 and n.t.=0.7 in HMR-1 and HMP-2 depleted embryos. Data from 5 different embryos and 12 time points per embryo. Ratios are significantly larger at all cell-cell contacts for embryos double depleted in HMR-1 and HMP-2, Wilcoxon rank-sum test. **E**. Tensions inferred from angles as a function of tension predicted from myosin fluorescence intensity using the relation *γ*^NMY2^ = *⍺*. *I*^NMY2^ + *β*, with α=0.751 pN/(μm.ua), β=325 pN/μm. Cell-medium surfaces are plotted in blue and cell-cell contacts in red. Each point corresponds to a given interface and time point and is averaged over 5 embryos. The correlation is measured through a Pearson coefficient ρ=0.468. The solid black line indicates a perfect correlation. **F**. Residual tension *γ*^residual^ = *γ*^infer^ − (*⍺*. *I*^NMY2^ + *β*) at cell-medium and cell-cell interfaces as a function of E-cadherin fluorescence intensity (HMR-1::GFP). Each surface is colour coded according to the inset. Each point corresponds to a given interface and time point and is averaged over 5 embryos. Cytoplasmic background fluorescence was removed from the HMR-1 signal. The fit corresponds to a Hill function of the form 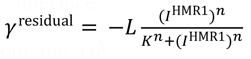 with a Hill coefficient n=1.78, a constant K=63.43 a.u. and a scale L=277 pN/μm. **G**. Surface tension at cell-medium interfaces (blue dots) and cell-cell interfaces (red dots) as a function of tension predicted from myosin and E-cadherin fluorescence intensity. Each point corresponds to a given interface and time point. Data is averaged over 5 embryos. The correlation is measured through a Pearson coefficient ρ=0.829. The solid black line indicates a perfect correlation.

### Cell-cell contact tension depends non-linearly on E-cadherin enrichment

The lower tensions observed at cell-cell interfaces compared to cortical interfaces suggested a role for intercellular adhesion proteins in the regulation of interfacial tension. Although previous work has highlighted roles for E-cadherin as well as L1CAM in *C. elegans* gastrulation ^47^, cells are primarily linked to one another via E-cadherin adhesion complexes at the 4-cell stage ^20^. Cadherins can decrease surface tension directly by increasing adhesion between cells and indirectly by decreasing myosin activity via downregulation of RhoA due to recruitment of p120-catenin by the cadherin cytoplasmic domain ^12,31^. Previous work in *Drosophila* suggested that cadherins contributed both directly and indirectly to tension at cell-cell contacts ^32^; whereas, in other organisms (such as mouse and zebrafish), cadherin-mediated adhesion only contributes indirectly through its regulation of myosin contractility at cell-cell contacts ^29^. We investigated how E-cadherin enrichment at intercellular contacts correlated with their surface tension.

First, we examined the temporal evolution of HMR-1 (E-cadherin) enrichment at intercellular contacts by acquiring time-lapses of embryos with a GFP fusion to the endogenous *hmr-1* gene, and processing the data in the same way as for NMY-2 (**Fig 6D-E, Fig S6A-D**). We then asked if there was a direct correlation between the evolution of tension at intercellular contacts and HMR-1 enrichment. At the onset of the 2-cell stage inside of the eggshell, the HMR-1 signal at the AB-P_1_ contact was initially visually indistinguishable from cytoplasmic background before gradually increasing until reaching a plateau mid-way through interphase for n.t.=0.5, when the contact became visible (**Fig S6A,C**). At the 4-cell stage, HMR-1 fluorescence intensity displayed marked differences, being largest at the contact between the AB cells and EMS, followed by ABa-ABp, ABp-P_2_ and weakest at the contacts with EMS-P_2_ (**Fig 6E, Fig S6B,D**). At contacts with P_2_, HMR-1 enrichment was initially visually indistinguishable from cytoplasmic background before steadily increasing throughout interphase, reaching a plateau by n.t.=0.5. Intriguingly, we found no obvious correlation between the tension at cell-cell contacts and their E-cadherin enrichment, suggesting that HMR-1 is not the only factor regulating cell-cell interfacial tension. We hypothesized that this may be because HMR-1 is interfaced to the F-actin cytoskeleton in some contacts but not in others. Therefore, we imaged the spatiotemporal localization of HMP-1 and HMP-2 (α- and β-catenin) that link E-cadherin to F-actin. However, their enrichment at contacts mirrored in all points that of HMR-1 (**Fig S6I-K**). Overall, this suggests a complex relationship between interfacial tension and cadherin localization.

Next, we investigated if tension at intercellular contacts could be predicted solely based on myosin enrichment, reasoning that any regulation emanating from E-cadherin signaling should be reflected in a change in levels of myosin at the cell-cell contact. First, we searched for evidence of the myosin downregulation at cell-cell contacts reported in previous studies ^31^. We reasoned that, if no regulation of myosin takes place, the myosin intensity at cell-cell contacts should be equal to the sum of the myosin intensity at the cellular cortices of the two cells in contact because the overall thickness of the cell-cell contact is similar to pixel size in our images. To quantify this, we computed the ratio of the myosin intensity at cell-cell contacts divided by the sum of the cortical myosin intensities at cortices (**Fig 6C-D, Fig S6D**). This ratio should be equal to 1 if no regulation takes place and less than 1 if downregulation occurs. We found that the ratio at cell-cell contacts was systematically lower than 1, apart from in the contacts ABp-P_2_ and EMS-P_2_ before n.t.=0.5, in agreement with the initial exclusion of P_2_ from the aggregate. Similar observations were made for p-MLC in 4-cell stage embryos outside the eggshell, where we found that this ratio was strictly lower than 1 in all cell-cell contacts except for EMS-P_2_ and that it correlated negatively with the fluorescence of HMR-1 (**Fig S6C**). Finally, we found that the myosin ratio was always significantly higher (from 2 to 5 fold depending on the contact) in double depletions of HMR-1 and HMP-2 than in controls (**Fig 6D**). Overall, this showed that myosin is downregulated at cell-cell contacts in the *C. elegans* embryo and that its downregulation is directly linked to high E-cadherin density.

Therefore, we first hypothesized that the tension at cell-cell contacts could be predicted from myosin intensity only, like for cell-mediums surfaces (**Fig 4E**), and plotted inferred tension at cell-cell contacts as a function of tensions predicted from myosin intensity measured at contacts *γ*^NMY2^ = *⍺*. *I*^NMY2^ + *β* (**Fig 6E, S6F**). This revealed that actual tensions at cell-cell contacts are systematically lower than those predicted from myosin enrichment alone, indicating that myosin alone is not sufficient to predict tension at cell-cell contacts. To determine if this was due to an additional direct contribution of E-cadherin binding across intercellular contacts, we computed a residual tension for each cell-cell contact measuring the contribution not attributable to myosin: 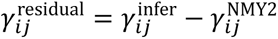 with 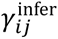 the tension between cells *i* and *i* inferred from contact angles and 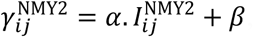, the tension predicted from myosin intensity only. When 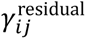 is plotted against the intensity of cadherin 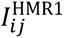 at the corresponding cell-cell contact (**Fig 6F, S6G**), we find that the residual tension is systematically negative and decreases non-linearly with increasing cadherin intensity (i.e. increases in absolute value) until reaching a plateau at the highest levels of cadherin enrichments. This relationship can be well fitted with a Hill function of the form 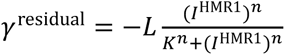 with L=277 pN/μm the maximal decrease of surface tension at infinite cadherin concentration, K=63.48 a.u. the HMR-1 intensity at which half of the maximal decrease occurred, *I*^HMR1^ the intensity of HMR-1 in a.u., and *n*=1.78 the Hill coefficient. This Hill coefficient larger than 1 suggests a possible cooperativity effect in the direct contribution of cadherin to cell-cell interfacial tensions. Interestingly, we observed a clustering of the points in the residual as a function of the nature of the contact (**Fig 6F**), mirroring the clear separation of contacts in terms of measured levels of HMR-1 (**Fig 6B**). From the previous findings, we decided to incorporate both contributions of cadherins and myosin to account for their combined empirical influence to contact tensions as follows 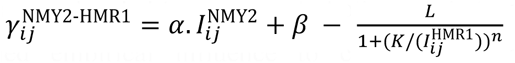. Doing so, we found that all surface tensions in the embryo could be well predicted based on myosin and cadherin intensities (**Fig 6G**), as judged by comparison to inferred values and the significant improvement in Pearson correlation score, from 0.47 to 0.83. In summary, we demonstrated that the tensions at cell-cell contacts are defined by both the level of myosin, whose enrichment is affected by cadherin-mediated signaling, and by the level of cadherins, through a non-linear direct negative contribution.

## Discussion

*Caenorhabditis elegans* is a rapidly developing organism that forms embryos with a fixed lineage specification and a characteristic cellular arrangement within an eggshell, making the characterization of cellular mechanical properties challenging. Here, we used quasi-static tension inference to find tensions that mechanically characterize the early *C. elegans* embryo. Our findings show that inferred relative tensions accurately and quantitatively predict the mean temporal evolution of cell shape and cellular arrangement in the embryo when used in foam-like simulations. This suggests that the arrangement of the embryo at the 2 and 4-cell stages can be relatively well approximated to a quasi-static foam. Combining quantitative imaging of cortical myosin fluorescence with tension inference, we could calibrate all surface tensions over time, including those at cell-cell contacts that cannot be measured directly. This approach enables the reconstruction of a spatiotemporal map of all tensions both inside and outside the eggshell, establishing a novel myosin-informed inference strategy for time-dependent force inference. Using AFM measurement on embryos outside of their eggshell, we furthermore establish a correlation between cortical myosin intensity and absolute values of cell-medium tensions, allowing to associate an absolute order of magnitude to our tension maps. Characterizing the spatiotemporal evolution of all surface tensions reveals the driving forces shaping the embryo and uncovers a clear direct contribution of cadherins to cell contact tension. This contribution manifests as a significant non-linear decrease in tension with increasing cadherin enrichment, which contrasts with previous reports on the negligible direct role of cadherins to contact tensions in mouse and zebrafish embryonic cells ^29,30^ or *Drosophila* somatic cells ^32^.

### Spatiotemporal mechanical changes in the early *C. elegans* embryo

Myosin-informed surface tension inference allowed us to draw a panorama of the spatiotemporal mechanics of the early *C. elegans* embryo and dissect the mechanical changes that drive cell arrangement. Compaction of ABa, ABp, and EMS in the 4-cell stage embryo was driven by a greater increase in cell-medium cortical tension in these cells compared to the tension at their cell-cell contacts. Conversely, P_2_ remained relatively separate from the EMS-ABx(2) aggregate because its absolute cortical tension remained lower than the tension at its contacts with EMS and, to a lesser extent, ABp. This was particularly apparent in embryos without an eggshell, in which the absolute cortical tension in P_2_ was ∼1.5 fold lower than in the EMS-P_2_ contact. At the molecular level, increases in cell-medium cortical tension were driven by myosin recruitment, while at intercellular contacts myosin enrichment was reduced, perhaps due to signalling downstream of E-cadherin or through action of the RhoGAP PAC1 ^48^. A more thorough understanding of the molecular mechanisms regulating surface tensions will represent an interesting future direction.

The low absolute cortical tension in P_2_ is intriguing as it is a type of stem cell and maintenance of pluripotency in mammalian *in vitro* stem cell culture relies on inhibition of contractility and a low surface tension ^8,49^. Therefore, as in mammalian systems, differentiation of cells in *C. elegans* may involve an increase in surface tension. Our experiments also revealed that regulation of cell-medium surface tension showed some differences across cell types. Indeed, although cortical myosin enrichment was a good overall predictor of the magnitude of cell-medium cortical tension, close examination of the temporal evolution of this relationship revealed differences between cell types (**Fig S4E**). For example, tension in P_2_ and EMS increased more steeply with myosin enrichment than AB and ABx. This may reflect differences in the structure ^43^ and proteic composition of the F-actin cortex ^50–52^ or differences in the phosphorylation state of myosin. A more precise understanding of the molecular control of surface tension will need to take the F-actin cortex structure into consideration. Comparison of our mechanical panorama to transcriptomic atlases of early *C. elegans* embryos ^53^ may help to understand how gene expression and cell mechanics interplay to shape developing embryos. Additionally, our direct measurements of cell-medium tension and cortical myosin intensity at cell surfaces revealed large variability between embryos (**Fig S3, S4**). This suggests that robust development does not require a tight control of absolute surface tensions or absolute myosin expression but rather a tight control over how myosin is partitioned and regulated between cells in each round of division. Further work will be necessary to test this hypothesis.

### Direct and indirect contributions of cadherins to tension at intercellular contacts

Cadherin-mediated adhesion can contribute to surface tension at intercellular contacts both directly and indirectly. While previous studies have mostly highlighted the prevailing indirect role of cadherins in regulating myosin activity ^29,30^, our findings challenge this notion. By examining cellular mechanical changes and molecular markers of contractility and adhesion, we found both significant direct and indirect contributions of cadherins to tension at cell contacts. Our results show that cadherin signaling can lead to a ∼50% decrease in myosin intensity at cell-cell contacts, although this decrease varies between cell-cell contacts (**Fig 6D, S6D**). At all cell-cell contacts, inferred tensions based on myosin fluorescence intensity were systematically larger than actual inferred tensions (**Fig 6E**), suggesting a direct cadherin contribution to reducing interfacial tension. Residual tension, calculated as the actual inferred tension minus the tension expected from myosin enrichment, decreased with increasing cadherin enrichment, saturating at higher cadherin intensities. This behavior suggests a cooperative binding mechanism, likely due to the cis-binding of cadherins ^54^. Notably, saturation at high cadherin enrichment is consistent with observations that HMR-1 overexpression does not cause phenotypic changes ^55^. Further research will be needed to determine if this saturation also affects cadherins’ indirect contributions via myosin activity regulation and to quantify the roles of other intercellular adhesion proteins in early *C. elegans* embryos ^47^. In addition, it will be particularly interesting to examine how the direct and indirect contributions of cadherin evolve with time in each intercellular contact and in particular in ABp-P2, which is the contact that forms the latest.

### Model limitations and future perspectives for mechanical inference

Through tension inference and simulation, we demonstrated that the mean shape of *C. elegans* embryos at the 2- and 4-cell stages can be well characterized by a static foam-like model (**Figs 2 and 5**). This result contrasts with modeling approaches relying on attractive and repulsive potentials between cells, leading to correct cell topologies, but not accurate shapes ^20–23^. Our static foam-like model may however not fully capture all dynamic cell shape events and transient cortical inhomogeneities observed in some cells within the embryo (see P_1_ close to the contact with AB in **Video S3** towards the end of the 2-cell stage). These rapid shape changes, such as cytokinesis, occur on typical cortical viscous relaxation timescales 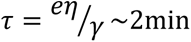 (with *η*∼10 Pa.s a typical viscosity^56^, *e*∼0.5μm a typical thickness ^57^, and *γ*∼500pN/μm a typical tension of the cortex). Recent research has shown that viscous surface resistance can lead to anisotropic tensions and that heterogeneities or gradients in surface contractility generate cortical flows ^10^. Both effects are sufficient to account computationally for complex dynamic shapes in 3D, such as those observed in cytokinetic cells ^56,58^, with great accuracy. Several models of interacting cells in 3D have been recently devised, incorporating viscous dissipation in various forms ^22,33,59,60^. However, none of these models explicitly account for the generic actomyosin depletion due to E-cadherin mediated signaling observed at cell-cell contacts in multiple species, including the *C. elegans* embryo (**Fig 6A-D**). Such depletion is expected to significantly reduce the viscous resistance to cell contact extensions. Instead, these models assume a single uniform cortical tension and a negative adhesive tension at contacts, similar to a passive non-living foam. Future modeling work will be necessary to accurately incorporate spatiotemporal inhomogeneities in cortical actomyosin based on experimental evidence. From an inverse problem perspective, novel methods must be devised to solve dynamic force inference problems that explicitly account for viscous surface flows, stresses, and a wider variety of force types. Recent work proposed a stochastic optimization approach to inference, allowing for the continuous capture of relative forces across divisions by including more complex forces than surface tensions ^61^. However, this approach still relies on the assumption of quasi-static cell shape evolutions, which may be questionable in contexts such as cytokinesis ^56^. Another study aimed to infer non-uniform and anisotropic tensions in pressurized epithelial layers but did not explicitly consider viscous stresses ^62^. Therefore, significant computational and experimental effort is still necessary to adapt force inference methods to viscous-active dynamics while precisely relating non-uniform and anisotropic active cortical stress to underlying protein activity.

## Conclusion

Overall, our study characterizes the dynamic spatiotemporal mechanical changes that shape the early *C. elegans* embryo, allowing us to predict cell arrangement from surface tensions at the 2- and 4-cell stage, and reveals the direct negative contribution of cadherins to interfacial tension.

## Author contributions

KY, HT, and GC conceived and designed the study. KY did most of the experiments and initial data analysis. SI, MP and HT designed and carried out the tension inference and numerical simulations and wrote the Supplemental Information. KY, SI, MP and FD performed the data analysis. JP and NG carried out control experiments. KY, SI, MP, HT and GC made the figures. KY, HT, and GC wrote the manuscript. All authors commented on the manuscript.

## Acknowledgements

We thank past and present members of the Charras and Turlier labs for comments and discussions. We thank Dr Grégoire Michaux (IGDR, Rennes, France) for discussions during the study and feedback on the manuscript. We thank anonymous reviewers for their constructive comments. We also thank Prof. Akatsuki Kimura in National Institute of Genetics for kind supports to KY. KY was supported by a Grant-in-Aid for JSPS Fellows (16J09469) and research grants from the Uehara Memorial Foundation. KY was supported by a European Research Council consolidator grant (CoG-647186) to GC. NWG is supported by the Francis Crick Institute that is supported by the Francis Crick Institute (N.W.G.), which receives its core funding from Cancer Research UK (FC001086), the UK Medical Research Council (FC001086), and the Wellcome Trust (FC001086). SI was supported by a PSL-UCL doctoral internship program and a Monge scholarship from Ecole Polytechnique. HT, SI, MP and FB are supported by the CNRS and Collège de France. HT has received funding from the European Research Council (ERC) under the European Union’s Horizon 2020 research and innovation program (Grant agreement No. 949267). Some *C. elegans* strains were provided by the CGC, which is funded by NIH Office of Research Infrastructure Programs (P40 OD010440). FL238 was kindly provided by Drs Anne Pacquelet and Grégoire Michaux Group in IGDR, Université de Rennes, France.

## Supplemental figures

**Figure S1:**
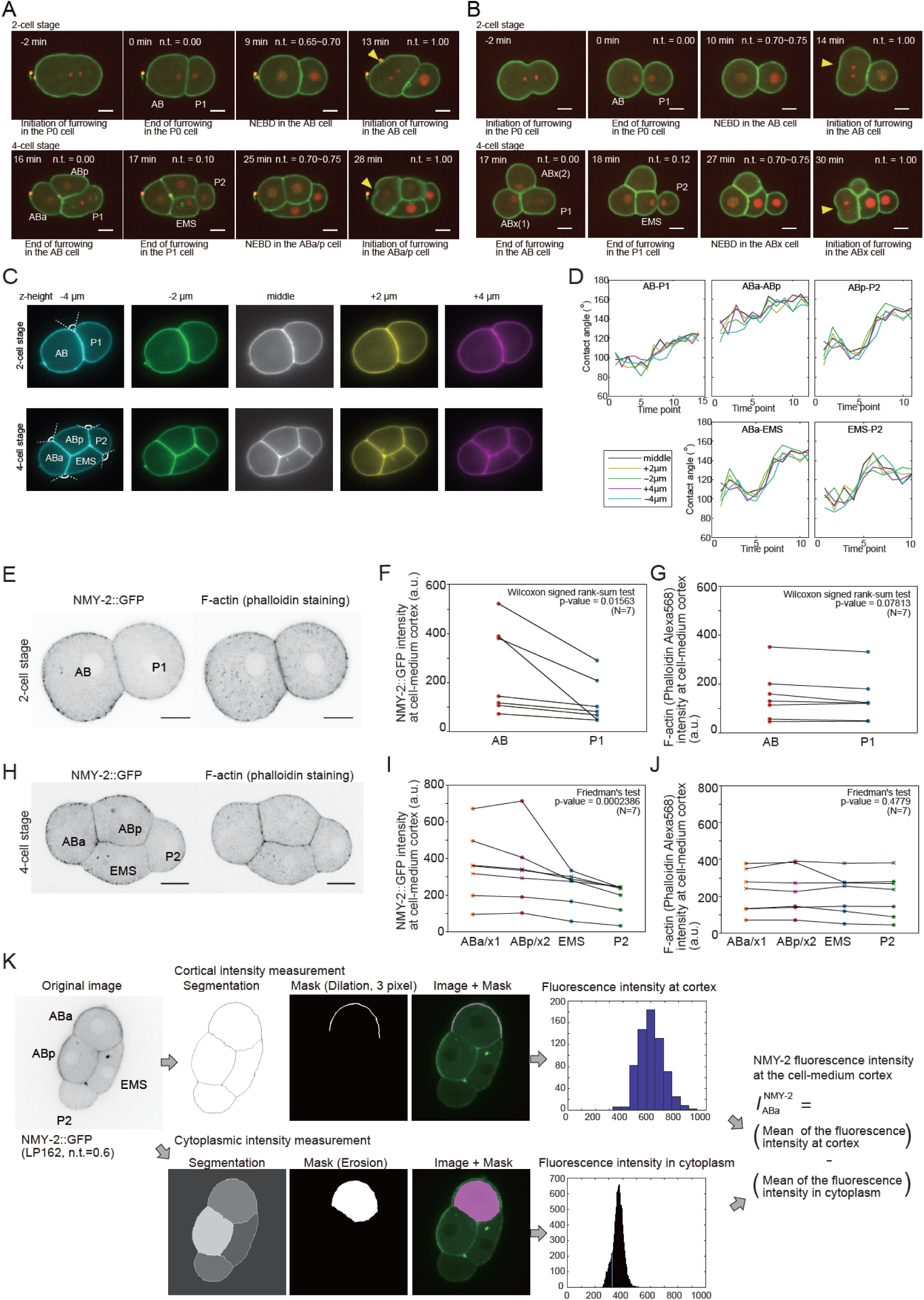
**(A-B)** Representative evolution of the cell arrangement in 2- and 4-cell *C. elegans* embryos. Cell membranes are visualized using GFP-PH-PLCδ and histones are visualized using mCherry::his-58. Scale bars =10μm. Yellow arrowheads indicate the position of furrowing. Cell names are indicated on each panel. Actual times are indicated in the top right corner. Normalised times (n.t.) are indicated in the top right corner. For the 2-cell stage, n.t.=0 was chosen as the time when P_0_ finishes its division and for the 4 cell-stage, it was chosen as the time when AB finishes its division. For the 2-cell stage, n.t.=1 was chosen as the time when AB starts to furrow and, for the 4-cell stage, it was chosen as the moment when ABa/p/x begins to furrow. **A**. Representative evolution of cell arrangement for *C. elegans* embryos in the eggshell. **B**. Representative evolution of cell arrangements for *C. elegans* embryo outside of the eggshell. **C**. Consecutive images of a confocal microscopy stack taken at different optical planes. The position of each plane is given relative to the mid-plane of the embryo. Top row: 2-cell stage. Bottom row: 4-cell stage. The name of each cell is indicated on the left most image. The angles of contact are displayed in the leftmost column. **D**. Temporal evolution of the angles of contact shown in C. Each line represents a different optical plane. Angular evolution is not affected by small differences in the optical plane chosen for measurement. **E**. NMY-2::GFP and F-actin distribution in the midplane of 2-cell stage *C elegans* embryos. The name of each cell is indicated on the images. Scale bar = 10μm. **F**. NMY-2 fluorescence intensities measured in the cortex of each cell. Data from 7 embryos, each embryo appears as a separate dot. Cells from the same embryo are linked by black lines. Fluorescence intensities were compared with a signed Wilcoxon rank-sum test. **G**. F-actin fluorescence intensities measured in the cortex of each cell. Data from 7 embryos, each embryo appears as a separate dot. Cells from the same embryo are linked by black lines. Fluorescence intensities were compared with a signed Wilcoxon rank-sum test. **H**. NMY-2::GFP and F-actin distribution in the midplane of 4-cell stage *C elegans* embryos. The name of each cell is indicated on the images. Scale bar = 10μm. **I.** Same as F but for the 4-cell stage. Fluorescence intensities were compared with a Friedman’s test. **J**. Same as G but for the 4-cell stage. Fluorescence intensities were compared with a Friedman’s test. **K**. Image analysis pipeline for extracting fluorescence intensities at interfaces. This is exemplified on NMY-2::GFP at the 4-cell stage. Top row shows cortical intensity measurement and the bottom row the cytoplasmic intensity measurement. For the cortical intensity, cell surfaces are segmented using Tissue analyzer. Then the mask for a specific surface (here ABa) is dilated to have a 3 pixel width. Then, it is convolved with the image to give a histogram of fluorescence intensities. For the cytoplasmic intensity, a mask of the cytoplasm is generated and eroded. This is then convolved with the image to generated a histogram of fluorescence intensities. Note that fluorescence intensities in the cortex are larger than in the cytoplasm. The NMY-2 fluorescence intensity at the cell-medium cortex is then computed as the difference between the means of the fluorescence at the cortex and the cytoplasmic intensity.

**Figure S2:**
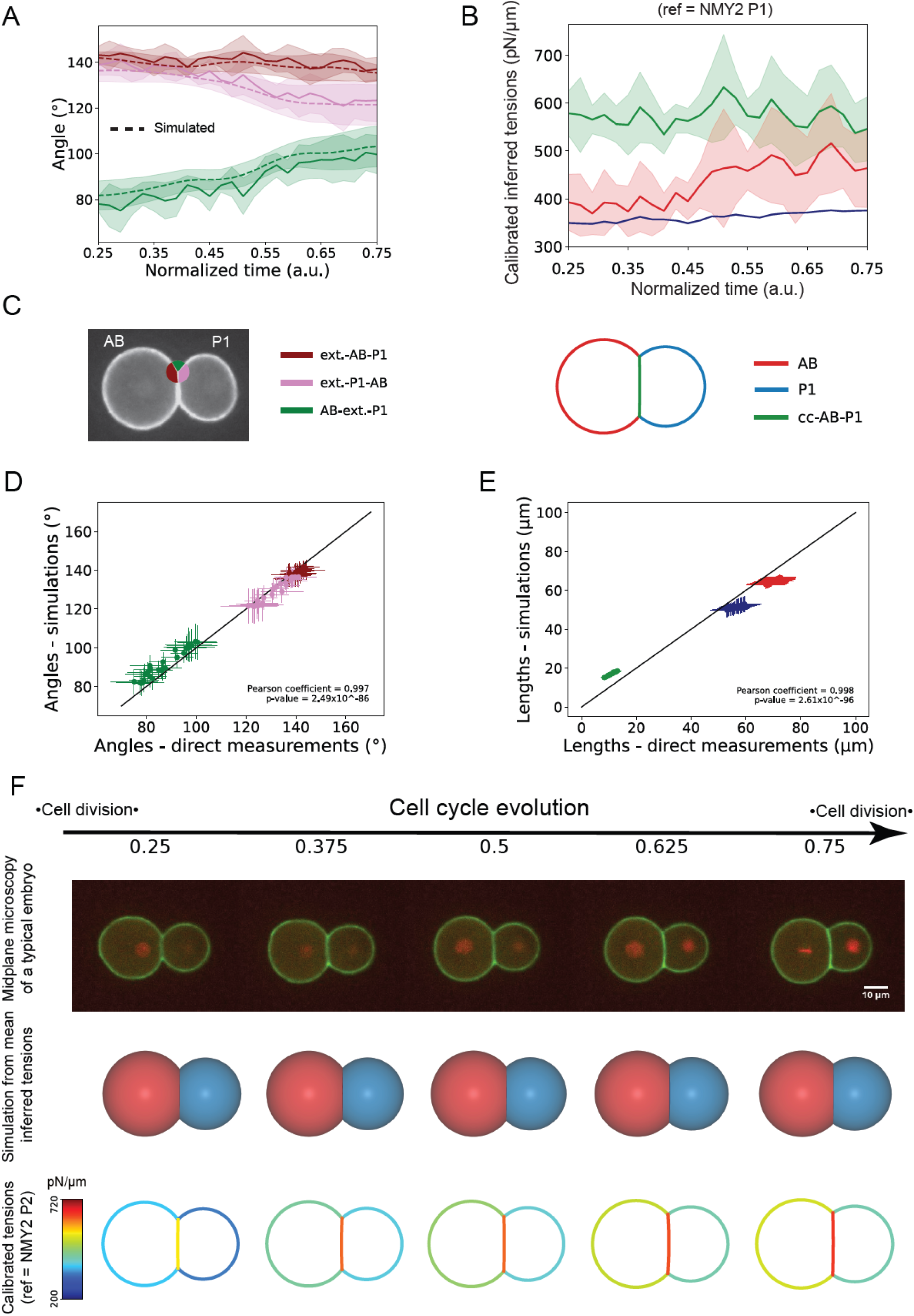
Inference and simulation of the 2-cell stage embryo outside of the eggshell. **A**. Temporal evolution of the external angles of contact at tricellular junctions. The solid line shows the mean and the shaded area represents the standard deviation (n=5 embryos). Time is normalised to the cell cycle and the color codes relating to external angles of contact are indicated in the panel C below. **B**. Temporal evolution of the mean inferred surface tensions calibrated in time and absolute value with the myosin fluorescence intensity in P1 and the affine parameters α=0.751 pN/(μm.ua), β=325 pN/μm. The solid line shows the average over 5 embryos and the shaded area represents the standard deviation. Time is normalised to the cell cycle. **C**. Left: Color code for contact angles in the 2-cell stage embryo used throughout the manuscript. Right: Color code for interfaces in the 2-cell stage embryo used throughout the manuscript. **D**. Plot of the contact angles obtained from simulations as a function of experimentally measured angles. Each data point represents one time point with standard deviation and is averaged over 5 embryos. The correlation is measured through a Pearson coefficient ρ=0.997. The black line shows the line of slope 1. **E.** Plot of the simulated lengths as function of experimentally measured lengths. Each data point represents one time point with standard deviation and is averaged over 5 embryos. The correlation is measured through a Pearson coefficient ρ=0.998. The black line shows the line of slope 1. **F.** Temporal evolution of inferred surface tensions allows prediction of the cell arrangement in embryos. First row: microscopy time series of a developing 2-cell stage embryo outside of the eggshell. The membrane is visualized with GFP-PH-PLCδ (green) and the histones are visualized with mCherry::his-58 (red). Scale bar=10µm. Second row: temporal evolution of the mean 3D embryo shape predicted by simulation (n=5 embryos). AB appears in orange and P_1_ in blue. Third row: temporal changes in surface tension in pN/μm. The tension in each surface is color coded with blue representing low tension and red high tension.

**Figure S3:**
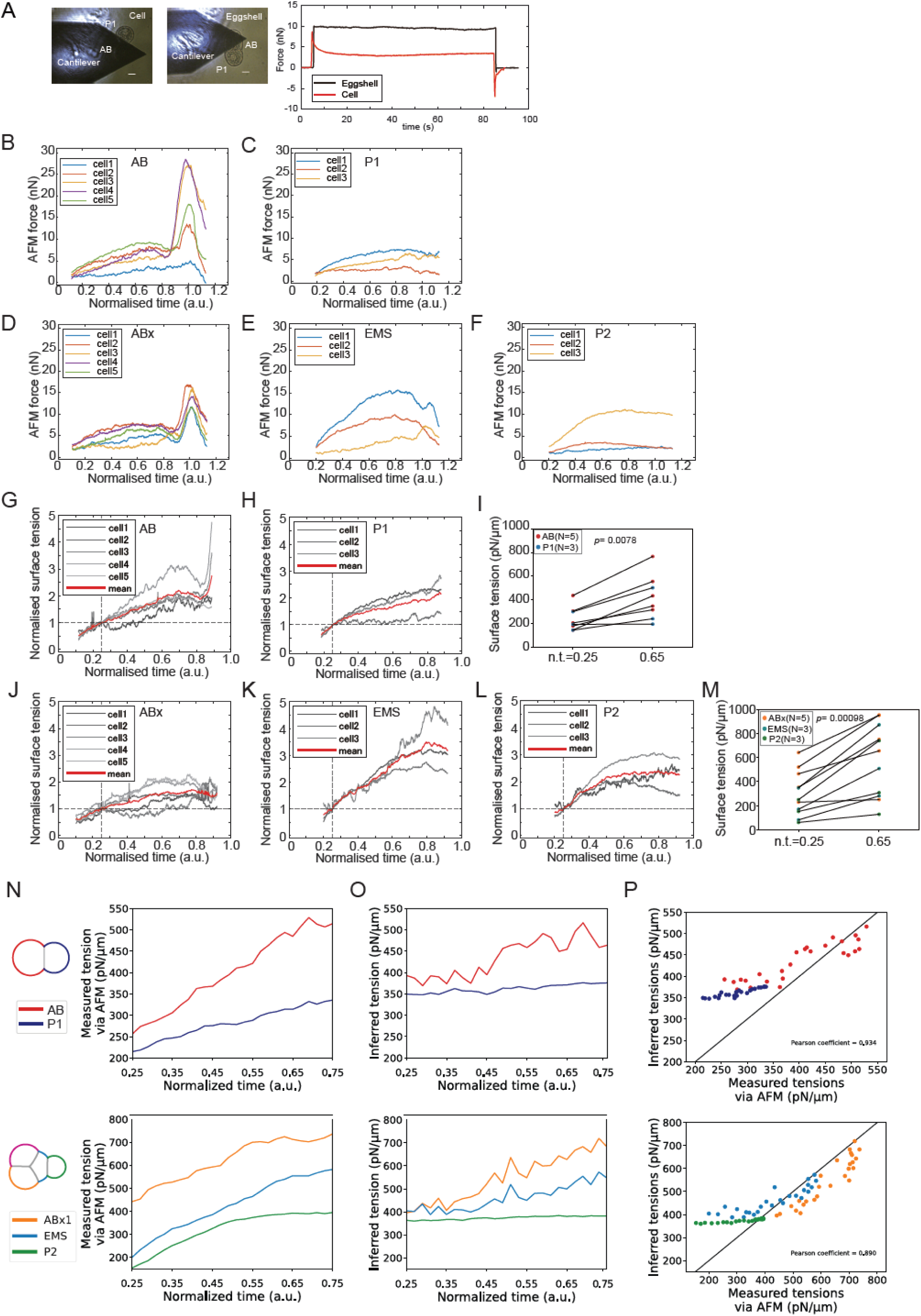
AFM measurements of cell surface tension. **A**. Representative AFM measurement of the mechanics of a cell and the eggshell. Left and middle: representative brightfield images of an AFM cantilever in contact with a 2-cell stage embryo without (left) and with (middle) an eggshell. The name of each cell is indicated on the images. Scale bar=10μm. Right: Representative graph showing the temporal evolution of the deflection force sensed by the AFM cantilever for a cell in an embryo without an eggshell (orange) or for the eggshell (blue). The cantilever is initially out of contact and senses no force. It is then brought into contact with the cell or the eggshell at t∼5s, leading to an increase in the measured force. The cantilever is then kept at a constant height for 80s before being retracted, leading to a return of the force to 0. While the cantilever is in contact, the force measured on the cell decreases due to viscoelastic relaxation stemming from biological processes such as cytoskeletal turnover. In contrast, no such relaxation is observed on the eggshell pointing to a solid-like behavior. **(B-F)** Temporal evolution of force measured by AFM when in contact with AB cells (B), P_1_ cells (C), ABx cells (D), EMS cells (E), and P_2_ cells (F). In all graphs the unit of force is in 10^−9^ N. Each line represents a different cell. Measurements are acquired at 5kHz and smoothed with a moving window of 1s. **(G-H, J-L)** Temporal evolution of cell-medium cortical tension normalised to the tension at n.t.=0.25 for AB cells (G), P_1_ cells (H), ABx cells (J), EMS cells (K), and P_2_ cells (L). Each grey line represents a different cell and the mean is plotted as a red line. (**I, M**) Tension in cells at n.t.=0.25 and n.t.=0.65. Lines link data points corresponding to the same cell. A Wilcoxon signed rank-sum test was used to compare tensions between time points. **N**. Temporal evolution of cell-medium cortical tensions measured by AFM for the 2-cell (top) and 4-cell (bottom) embryos outside of their eggshell. Each cell-medium interface is colour-coded as shown on the sketches to the left. **O**. Temporal evolution of cell-medium cortical tensions predicted by our myosin-informed inference approach for the 2-cell (top) and 4-cell (bottom) embryos outside of their eggshell. Colour codes are the same as in L. **P**. Inferred cell-medium cortical tension as a function of the cortical tension measured by AFM for the 2-cell (top) and 4-cell (bottom) embryos outside of their eggshell. Colour codes are the same as in L. (**N-O**) AFM data is from the cells in B-F and fluorescence data is averaged over 5 embryos.

**Figure S4 related to Figure 4:**
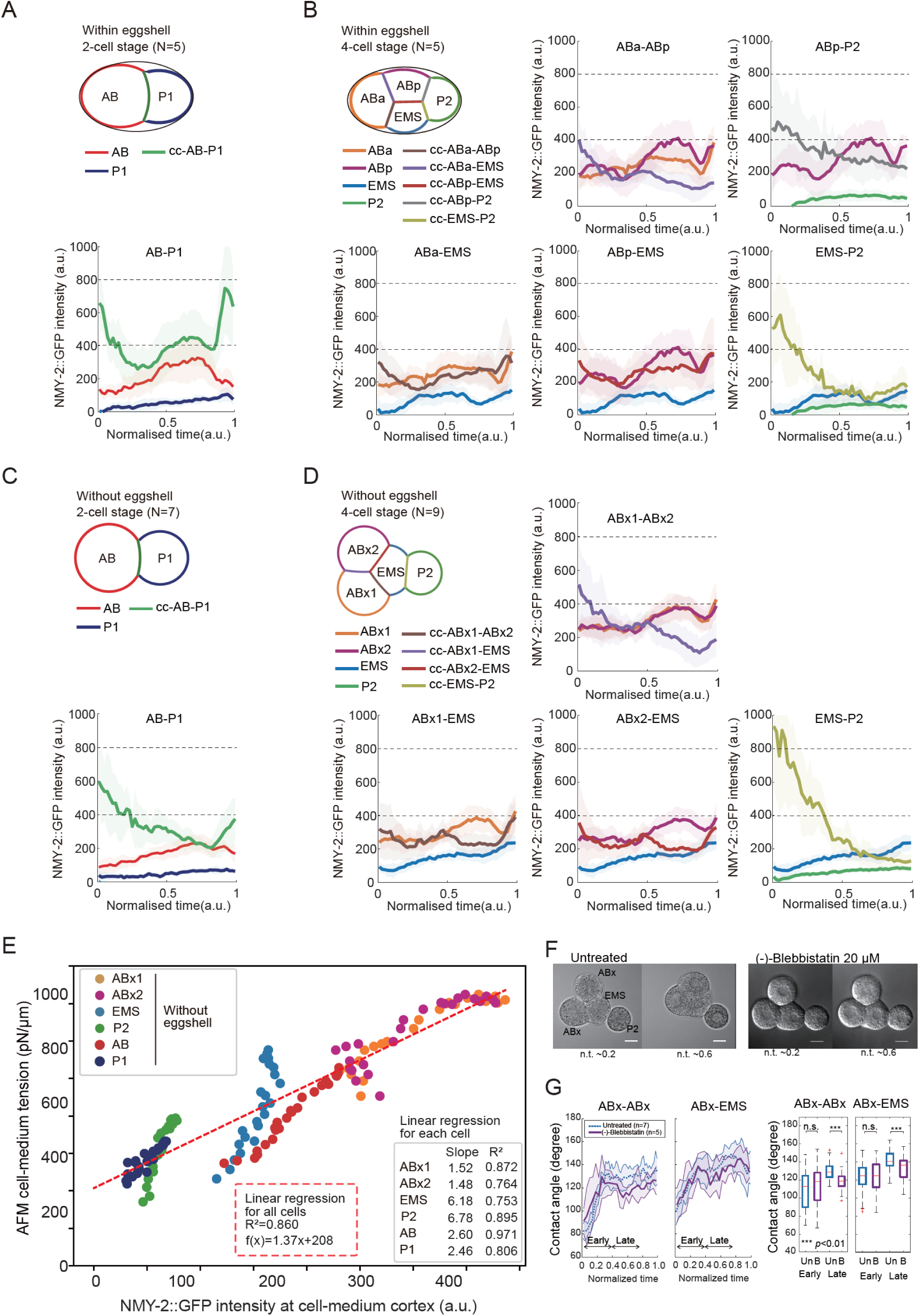
Temporal evolution of NMY-2 in the surfaces of the 2- and 4-cell *C. elegans* embryos with and without eggshell. **(A-D)** The pictograms indicate the color code for the surfaces of the embryos. In all graphs, the solid line is the average and the shaded region depicts the standard deviation. N≥5 embryos for each condition. Each graph shows the evolution of the fluorescence intensity of myosin in an intercellular contact (indicated as cc) and in the cortices on either side of that contact. All intensities are given in a.u. **A.** Temporal evolution of NMY-2 fluorescence intensity in 2-cell stage embryos within the eggshell. **B.** Temporal evolution of NMY-2 fluorescence intensity in 4-cell stage embryos within the eggshell. **C.** Temporal evolution of NMY-2 fluorescence intensity in 2-cell stage embryos outside of the eggshell. **D.** Temporal evolution of NMY-2 fluorescence intensity in 4-cell stage embryos outside of the eggshell. **E**. Cortical tension measured by AFM is plotted as a function of cell–medium myosin fluorescence intensity. Each data point corresponds to a single time point and represents the average over 5 embryos, with standard deviation indicated. The dashed red line indicates the best linear regression to the data from all cells (R^2^=0.86), with slope and intercepts α=1.37 pN/(μm.ua), β=208 pN/μm. The slopes of the regression for each cell type are given in inset. **F**. Bright field images of 4-cell stage *C. elegans* embryo without an eggshell untreated (left) and blebbistatin treated (right) at n.t.=0.2 and 0.6. Scale bar 10 μm. **G**. Left : Temporal evolution of contact angles between ABx-ABx and ABx-EMS in untreated (blue) and blebbistatin treated conditions (purple). Right : Comparison of time-averaged contact angles between ABx-ABx and ABx-EMS in untreated (blue, “Un”) and blebbistatin treated condition (purple, “B”) at early (n.t.= from 0 to 0.4) and late (n.t.= from 0.4 to 0.8) stages. Contact angles were compared with a Wilcoxon rank-sum test.

**Figure S5:**
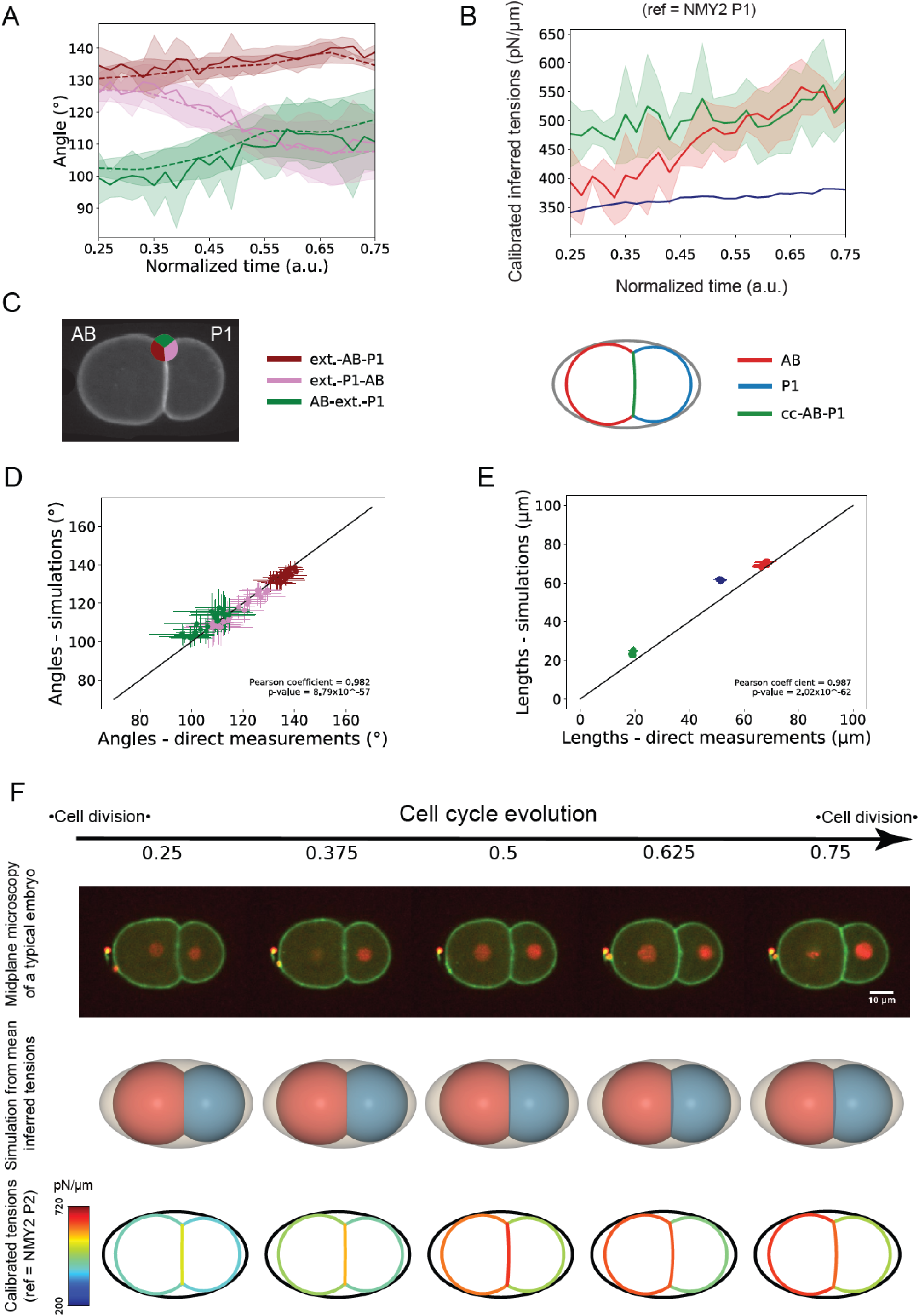
Simulation of the 2-cell stage embryo inside the eggshell. **A**. Temporal evolution of the external angles of contact at tricellular junctions. The solid line shows the mean and the shaded area represents the standard deviation (n=5 embryos). Time is normalised to the cell cycle and the color codes relating to external angles of contact are indicated in the panel C below. **B**. Temporal evolution of the mean inferred surface tensions calibrated in time and absolute value with the myosin fluorescence intensity in P1 and the affine parameters α=0.751 pN/(μm.ua), β=325 pN/μm. The solid line shows the average over 5 embryos and the shaded area represents the standard deviation. Time is normalised to the cell cycle. **C**. Left: Color code for contact angles in the 2-cell stage embryo used throughout the manuscript. Right: Color code for interfaces in the 2-cell stage embryo used throughout the manuscript. **D**. Plot of the contact angles obtained from simulations as a function of experimentally measured angles. Each data point represents one time point with standard deviation and is averaged over 5 embryos. The correlation is measured through a Pearson coefficient ρ=0.982. The black line shows the line of slope 1. **E.** Plot of the simulated lengths as function of experimentally measured lengths. Each data point represents one time point with standard deviation and is averaged over 5 embryos. The correlation is measured through a Pearson coefficient ρ=0.987. The black line shows the line of slope 1. **F.** Temporal evolution of inferred surface tensions allows prediction of the cell arrangement in embryos. First row: microscopy time series of a developing 2-cell stage embryo outside of the eggshell. The membrane is visualized with GFP-PH-PLCδ (green) and the histones are visualized with mCherry::his-58 (red). Scale bar=10µm. Second row: temporal evolution of the mean 3D embryo shape predicted by simulation (n=5 embryos). AB appears in orange and P_1_ in blue. The shell is indicated in grey. Third row: temporal changes in surface tension in pN/μm. The tension in each surface is color coded with blue representing low tension and red high tension. The shell is indicated in black.

**Figure S6:**
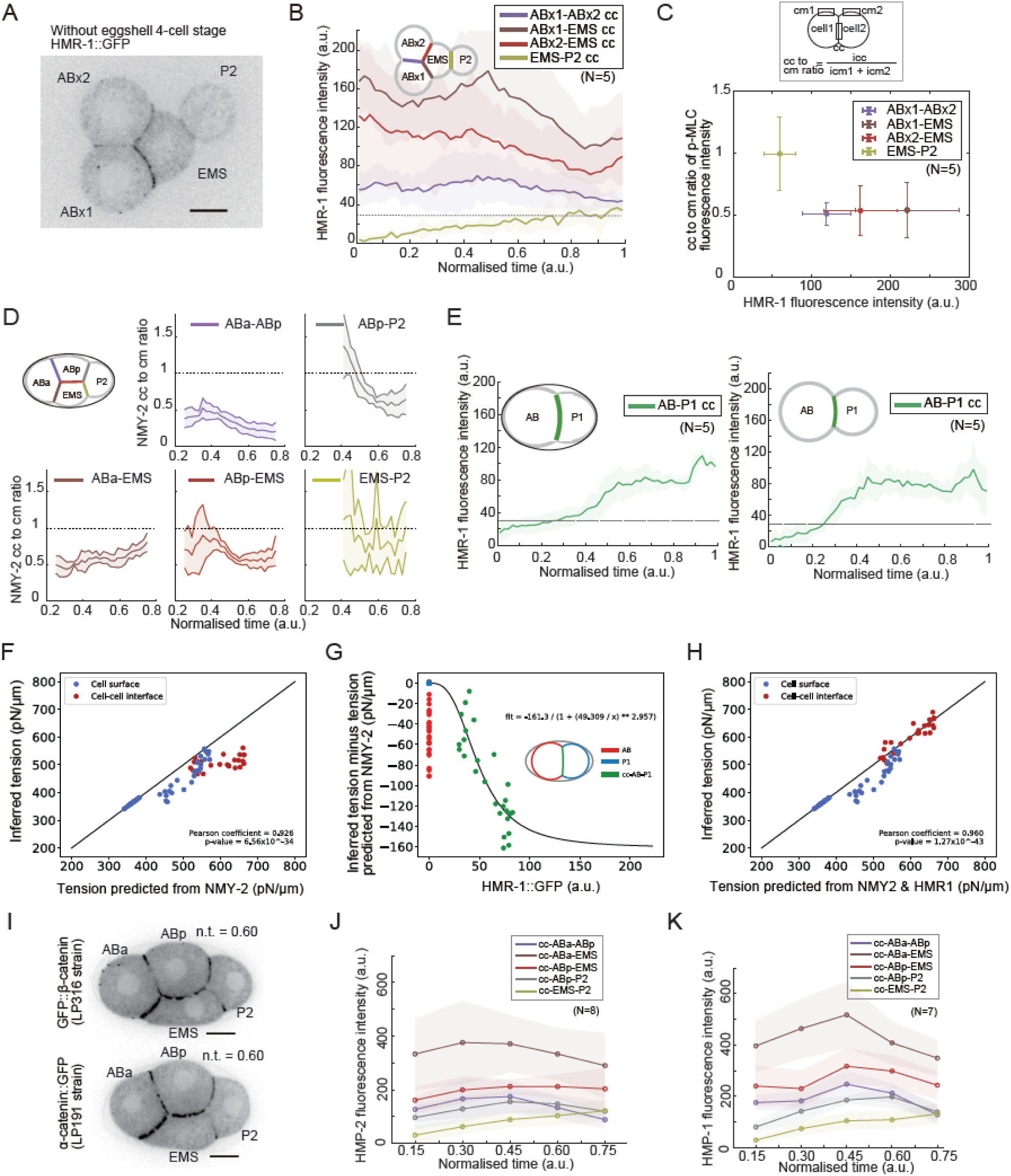
Temporal evolution of HMR-1, α-catenin, β-catenin and myosin at cell-cell contacts in the *C. elegans* embryos. A. E-cadherin-GFP (HMR-1) localization in a representative embryo at the 4-cell stage without the eggshell. Scale bar=10μm. The identity of each cell is indicated on the image. B. **B**. Temporal evolution of HMR-1 fluorescence intensity in 4-cell stage embryos without the eggshell. The inset pictogram indicates the position and colour-code of cell-cell contacts in the embryo. The solid line is the average and the shaded region depicts the standard deviation. N=5 embryos for each condition. The dashed line indicates the threshold fluorescence below which HMR-1 enrichment at cell-cell contacts is not visually distinguishable from background. C. Top: sketch of the measurement of the ratio of p-MLC fluorescence intensity at intercellular contacts and at the cell-medium cortex in the two cells in contact. The p-MLC data is taken from Fig 4B-C. The ratio is computed as the intensity i_cc_ in the cell-cell contact divided by the sum of the intensities i_cm1_ and i_cm2_ in the cortices. Bottom: The ratio of myosin at cell-cell contacts to the myosin at cortices is plotted as a function of HMR-1 enrichment in each cell-cell contact (taken from the measurements in D, top panel is a representative image). Whiskers indicate the standard deviation. Data is averaged over 5 embryos. p-MLC decreases with increasing HMR-1 in the contact. D. Comparison between NMY-2::GFP fluorescence intensity at intercellular contacts and at the cell-medium cortex in each pair of cells in contact. The ratio is computed as the intensity I_cc_ in the cell-cell contact divided by the sum of the intensities I_cm1_ and I_cm2_ in the cortices. Data from 5 different embryos and 18 or 13 time points per embryo. **E. Left :** Temporal evolution of HMR-1 fluorescence intensity at the AB-P_1_ contact for an embryo with an eggshell. Right : Temporal evolution of HMR-1 fluorescence intensity at the cell-cell contacts in the 4-cell stage embryo with an eggshell. **F**. Tensions inferred from angles as a function of tension predicted from myosin fluorescence intensity in 2-cell embryos within the eggshell. Tensions were inferred from myosin intensity using the relation *γ*^NMY2^ = *⍺*. *I*^NMY2^ + *β*, with α=0.751 pN/(μm.ua), β=325 pN/μm. Cell-medium surfaces are plotted in blue and cell-cell contacts in red. Each point corresponds to a given interface and time point and is averaged over 5 embryos. The correlation is measured through a Pearson coefficient ρ=0.926. The solid black line indicates a perfect correlation. **G**. Residual tension *γ*^residual^ = *γ*^infer^ − (*⍺*. *I*^NMY2^ + *β*) at cell-medium and cell-cell interfaces in 2-cell stage embryos within the eggshell as a function of E-cadherin fluorescence intensity (HMR-1::GFP). Each given cell-cell interface is attributed a different color. Each point corresponds to a given interface and time point and is averaged over 5 embryos. Cytoplasmic background fluorescence was removed from the HMR-1 signal. The fit corresponds to a Hill function of the form 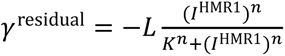 with a Hill coefficient n=2.96, a constant K=49.31 a.u. and a scale L=161.3 pN/μm. **H**. Surface tension at cell-medium interface (blue dots) and cell-cell interfaces (red dots) as a function of tension predicted from myosin and E-cadherin fluorescence intensity in 2-cell stage embryos within the eggshell. Each point corresponds to a given interface and time point. Data is averaged over 5 embryos. The correlation is measured through a Pearson coefficient ρ=0.980. The solid black line indicates a perfect correlation. **I**. Top: Representative images of β-catenin (HMP-2) in the 4-cell *C. elegans* embryo with an eggshell. Bottom: Representative images of α-catenin (HMP-1) in the 4-cell *C. elegans* embryo with an eggshell. Scale bars=10µm. **J**. Temporal evolution HMP-2 fluorescence intensity at the cell-cell contacts in the 4-cell stage embryo with an eggshell. The solid line is the average and the shaded region depicts the standard deviation. Data from N=8 embryos. **K**. Temporal evolution HMP-1 fluorescence intensity at the cell-cell contacts in the 4-cell stage embryo with an eggshell. The solid line is the average and the shaded region depicts the standard deviation. Data from N=7 embryos.

## STAR Methods

### C. elegans strains

The *C. elegans* strains used for this study are listed in Table S1. *C elegans* were maintained using a standard procedure for *C. elegans* culture ^63,64^. Briefly, worms were kept on NGM agar plates seeded with OP50 E. coli at 20C until use. *C. elegans* embryos were dissected from adult worms and collected in 0.75× egg salt (118 mM NaCl, 40 mM KCl, 3.4 mM CaCl2, 3.4 mM MgCl2, 5 mM HEPES pH 7.2) under the stereoscopic microscope.

### RNA interference

RNA interference for *hmr-1*, *hmp-2* was done by the standard feeding method ^65^. The C. elegans RNAi library (Source BioScience, Nottingham, UK) was used ^66^ for *hmp-2*. For *hmr-1* RNAi, the E. coli strain described previously was used ^20^. Briefly, E-coli were seeded on NGM-IPTG plate and they were kept at room temperature overnight before use. *C. elegans* larvae at L3∼L4 stage were plated on the plates and kept at 20C for 30 to 36 hours before dissection.

### Eggshell removal

Eggshell was removed using a previously described method with modification ^67^. Briefly, gravid worms were dissected in 0.75x Egg salt and 1-cell or 2-cell stage embryos were collected using mouth-pipette. The embryos were treated in bleach mix for 90 seconds. Then the embryos were washed in Shelton’s growth medium (SGM) ^68^ three times. Lastly, a fine glass capillary (GD-1; Narishige, Tokyo, Japan) was pulled on a Sutter Flaming brown pipette puller. The embryos were imaging on a stereoscopic microscope and the glass capillary was used to remove the eggshell with mouth pipetting. The cells were then kept in SGM before use at room temperature.

### Atomic Force Microscopy

For cell-medium cortical tension measurements, firstly the eggshell was removed at the 1-cell stage or 2-cell stage. Next, the embryos were transferred to a glass bottom Petri dish (WPI, Sarasota, FL, USA) filled with 0.75x Egg salt with a drop of SGM. To keep the temperature of the medium below 25C, a custom-made temperature cooler was employed. Tension measurements were performed with a CellHesion atomic force microscope (JPK instruments, Berlin, Germany) mounted on an IX81 inverted microscope (Olympus, Japan) equipped with a FV1000 confocal laser scanner unit. Flat cantilevers with a nominal spring constant of k=0.03 N/m (Arrow TL1Au, Nanoworld, Neuchâtel, Switzerland) were used for measurements. The sensitivity was calibrated by acquiring a force curve on a glass coverslip and the spring constant was calibrated using the thermal noise calibration method implemented in the AFM controller software. With these experimental conditions, we estimate the force resolution of AFM to be ∼200 pN. Two types of cortical tension measurements were performed: short-term compression experiments were used to measure cortical tension at a given time point and long-term compression experiments to follow the temporal evolution of cortical tension.

In short-term compression measurements, a force curve was acquired on the glass surface next to the cell of interest to allow for determination of cell height. Next, the cantilever was positioned above the cell of interest, the cantilever was approached towards to cell at a speed of 5 μm/s with a force setpoint of 10 nN. On average, cell height was reduced by 20 to 30% by application of force. After contact, force relaxed rapidly before reaching a plateau after ∼30s, consistent with previous reports on mammalian cells ^41^. The mean value of the force between 30-70 s after contact was used as input to calculate cell-medium cortical tension. After 80s contact, the cantilever was retracted. During these measurements, data was sampled at 5kHz and smoothed with a 1s window during postprocessing and the baseline of the data was adjusted so that the force value at the beginning and the end point was zero. The maximum cell radius during compression was obtained from a bright field image. The height of the cell before and during compression was measured following the method in ^45^. For static AFM measurements, we collected data from a total of 84 embryos, performing compression experiments on 153 cells. From each cell, we collected only one experimental force-compression curve. We verified that each of the cell compression curves we acquired reached a plateau after 30s, signifying that the cell does appears to have a clear surface tension. Measurements that did not reach a plateau were excluded from the later analysis and this led to the removal of approximately 5% of the experimental curves.

Long-term compression experiments used the same experimental protocol, except that the cantilever remained in contact with the cell until division. In these experiments, EG4601 embryos expressing GFP::H2B were imaged with the confocal microscope (a FV1000 laser scanning confocal head mounted on an IX81 microscope) to monitor the evolution of the cell cycle during compression. A force setpoint of 5 nN was chosen. After reaching this setpoint, the height of the cantilever chip was kept constant and we monitored its deflection, sampling at 5kHz. After converting to force, this data was smoothed with a 1s window during postprocessing and the baseline of the data was adjusted so that the force value at the beginning and the end point was zero. Drifts in laser position on the photodetector due to small temperature drift were corrected by measuring the laser position immediately before contact with the cell and immediately after retracting the cantilever and by applying a linear correction. The maximum cell radius and cell height were measured as described above to calculate cell-medium cortical tension.

### Cortical tension calculation

The full derivation of the formula to calculate the cortical tension is detailed in the SI. Here, we do not make any geometric approximation for the shape of the profile ^41^, but we solve for the true Laplace’s profile, following ^42^. The cell is characterized by a cortical tension *γ* and is compressed by an AFM cantilever with a height *h* with a force *F*. We suppose that the adhesion of the cell to the substrate and to the AFM cantilever are negligible, such that the contact angle *θ* = 0. The maximum cell radius at the midplane parallel to the AFM cantilever is measured optically (see above) and denoted by *r*_1_. The free radius of the cell, measured before compression is denoted by *r*_0_.

The tension is given by force balance at the midplane and reads:

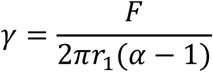

where *⍺* is a dimensionless parameter, which is determined using the AFM height *h* and the planar radius in the midplane *r*_1_ (see **Supplemental theory**).

### Live cell imaging

A spinning disk confocal microscope was used to acquire time-lapse images for LP162, LP172 and FL238 strains. For embryos with an eggshell, 1-cell stage embryos were placed in 0.75x egg salt on a 1% agar gel pad and this was gently covered with a coverslip. For embryos whose eggshell had been removed, embryos were placed in SGM on the coverslip, this was then covered by another coverslip and sealed with valap (1:1:1 mixture of petroleum jelly:lanolin:paraffin wax) to prevent drying. Embryos were imaged at room temperature with a spinning-disk head (CSU-10, Yokogawa, Japan) mounted on an IX71 inverted microscope (Olympus) equipped with a 60x oil objective (UPlanSApo, NA 1.35, Olympus) or 40x silicone objective (UPlanSApo, NA 1.25, Olympus). Images were acquired with an sCMOS camera (Prime 95B, Teledyne Imaging, Tucson, Az, USA) controlled by μManager ^69^. GFP and mCherry were excited with a 488nm and a 561nm laser, respectively. The exposure time was set such that the signal intensity was not saturated at any position.

### F-actin staining and immunostaining for p-MLC

*C. elegans* embryos were fixed after removing the eggshell. For phalloidin staining, the embryos were fixed for 30 minutes at room temperature in a solution containing 2% PFA, 0.25% Glutaraldehyde, HEPES (pH6.9) 100mM, EGTA (pH7.0) 50mM, MgSO_4_ 10mM, Triton X-100 0.2%, Dextrose 100mM after the eggshell removal. The buffer composition followed previous publications ^9,12^. After fixation, the embryos were washed in PBS three times, and then incubated with Phalloidin Alexa568 (Thermo Fisher scientific) for two hours at room temperature.

For immunostaining, the eggshell was removed from the embryos and these were fixed in 4% PFA, HEPES (pH6.9) 100mM, EGTA (pH7.0) 50mM, MgSO_4_ 10mM, Triton X-100 0.2%, Dextrose 100mM for 30 minutes at room temperature. The embryos were then washed with 0.1% Triton in PBS three times. They were then incubated with a rabbit anti-phosphomyosin S19 antibody (1:200 dilution, #3671, Cell signaling technology, Danvers, Ma, USA) diluted in PBS with horse serum and 0.01% Triton at 4C overnight. The fixed embryos were then washed with PBS three times. They were then incubated with goat anti-Rabbit Alexa568 in PBS at room temperature for two hours. The embryos were then washed with PBS three times.

The embryos were then placed on a coverslip coated with Poly-lysine to bind them to the surface. Images were acquired using a FV1200 laser scanning confocal head mounted on an IX83 microscope equipped with GaAsP detectors (Olympus). Optical sections were acquired at 0.25μm intervals along the z-axis.

### Image analysis

#### 1. Angle measurement

Images were semi-automatically segmented based on membrane fluorescence signal from PH-PLδ-GFP. Cell contours were obtained by thresholding of the intensity, and processed using a custom mesh generation pipeline based on distance transforms and Delaunay-Triangulation to generate a triangular tilling of the images and line contour segmentation based on the triangles of the tilling. Our method returns the borders of the cells as multi-material polygonal line, that can be used to extract all the junctional angles (see **Supplemental Theory**). We confirmed that angle measurement was not affected by small offsets in the plane of imaging from the embryo midplane (**Fig S1C,D**).

#### 2. Fluorescence intensity measurement

To measure average value of the fluorescent intensity along different parts of the cell contour, images were semi-automatically segmented using a Fiji plugin, TissueAnalyzer ^70^ (**Fig S1K**). The segmented images were subdivided into cortical regions, intercellular adhesions, and cytoplasmic regions. Cortical and cell-cell contact regions were then dilated to increase their width. Cytoplasmic regions were eroded to decrease their area and ensure no overlap with interfacial regions. Those regions were used to define ROI to obtain pixel values contained within each region (**Fig S1K**). A MATLAB script was written to extract pixel values, calculate a mean intensity for each ROI, as well as output histograms of fluorescence intensities.

We computed the fluorescence intensity in cell surfaces by subtracting the cytoplasmic fluorescence intensity from the cortical or cell-cell contact fluorescence intensity. Indeed, fluorescence in the cortex has two contributions: one from the cytosolic fraction and one from the fraction bound to the cortex ^71,72^. This is because the cortex is a loose actin mesh interpenetrated by cytosol and through which unbound proteins can diffuse. As tension generation only arises from myosins bound to the actin cortex, we subtract the cytoplasmic signal from the cortical signal to account for unbound myosins diffusing through the cortex.

### Statistical analysis

To confirm normality, the Shapiro–Wilk test was used. To confirm homoscedasticity, an F-test was used. If both normality and homoscedasticity were confirmed, Student’s t-test was used to compare means; In other cases, Wilcoxon’s rank sum test was used to compare median values between independent group and Wilcoxon’s signed-rank sum test was used for paired group. P<0.05 was considered to represent statistical significance. For these analyses, R (www.r-project.org) or MATLAB was used.

### Myosin-informed tension inference

We first infer ratios of surface tensions from angle measurements by solving a linear system of modified Young-Dupré equations, complemented by a constraint of mean tension equal to unity (see details in the **Supplemental theory**). Then we fix the scales of the tensions in space and in time using the evolution of myosin intensity values at the interface between P_2_ and the external medium. Finally, the regression between myosin intensity and tension values measured by AFM allows us to create spatiotemporal maps of absolute values of surface tensions within the embryo using P_2_ myosin intensity as temporal reference.

### 3D heterogeneous foam simulations and 2D sections

Knowing cells volumes and relative surface tensions (through inference), we minimize the surface energy *E* = ∑_{*ij*}_ *A*_*ij*_*γ*_*ij*_ (where *A*_*ij*_ stands for the interface between cells *i*, *j*, 0 being the cell medium by convention) under the constraints of conservation of the volume of each individual cell. We start from an initial triangular mesh with a topology similar to the corresponding wild-type embryo at the same stage. Using a conjugate-gradient numerical scheme to optimize the energy and a projection method to implement volume conservation constraints (see **Supplemental theory**), the mesh is evolved until a convergence criterium is reached, while remeshing operations are performed during optimization when necessary (when a topology change may occur for instance, or when the mesh is too distorted). A penalization energy (see **Supplemental theory**) is added to mimic the confinement within the shell (**Supplemental theory Fig 5**), whose shape is approximated to a prolate ellipsoid of dimensions given in **Supplemental theory Fig 4**. To mimic optical sectioning, we intersect the triangular mesh with a plane and we determine polygonal line contours of cells in the corresponding focal plane using the CGal library (https://www.cgal.org, see **Supplemental theory Fig 6**). Interface lengths can be easily calculated on this polygonal 2D representation of cells interface contours and compared to those obtained from segmentation meshes.

### Data availability

Raw experimental data, single embryo inference results and python notebooks used to generate figures are available from the UCL data repository (https://rdr.ucl.ac.uk/) with a unique doi (https://doi.org/10.5522/04/21977771).

Segmentation and force inference software are available on the following repository: https://github.com/VirtualEmbryo/foambryo2D

## Supplemental videos

**Video S1**. Time-lapse movie of a developing *C. elegans* embryo expressing a GFP-tagged membrane marker and mCherry-tagged histone marker and manually deprived of its eggshell from zygote stage. Embryos were observed from 1-cell to 2-cell stage, related to Figure S1, S3 (left), and from 2-cell to 4-cell stage, related to Figures 1, S1 (right). Scale bar = 10μm.

**Video S2**. Movies of 3D heterogeneous foam simulations of the *C. elegans* embryo without an eggshell at the 2-cell stage (left) and 4-cell stage (right), related to Figures 2, S2.

**Video S3**. Measurement of cortical tension dynamics using AFM, related to Figure 3, S3. The AFM cantilever was left in contact with the cell for the duration of its cell cycle.

**Video S4**. Time-lapse movie of a developing *C. elegans* embryo expressing a GFP-tagged membrane marker and mCherry-tagged histone marker within its native eggshell. Embryos were observed from 1-cell to 8-cell stage, related to Figures 2, 5, S1, S2, S4 (left). *C. elegans* embryo expressing a GFP fusion to the endogenous NMY-2 gene within the eggshell from 1-cell to 8-cell stage, related to Figures 4, S4 (right). Scale bar = 10μm.

**Video S5**. Movies of 3D heterogeneous foam simulations of the *C. elegans* embryo within the eggshell at the 2-cell stage (left) and 4-cell stage (right), related to Figures 5, S5.

**Video S6**. Time-lapse movie of a developing *C. elegans* embryo expressing a GFP fusion to the endogenous HMR-1 gene within the eggshell from 1-cell to 8-cell stage, related to Figures 6, S6. Scale bar = 10μm.

